# Successful Cardiac Resynchronization Therapy Reduces Negative Septal Work in Patient-Specific Models of Dyssynchronous Heart Failure

**DOI:** 10.1101/2024.05.13.593804

**Authors:** Amanda Craine, Adarsh Krishnamurthy, Christopher T. Villongco, Kevin Vincent, David E. Krummen, Sanjiv M. Narayan, Roy C. P. Kerckhoffs, Jeffrey H. Omens, Francisco Contijoch, Andrew D. McCulloch

**Author notes:** Contributed equally (Amanda Craine, Adarsh Krishnamurthy, Francisco Contijoch).

## Abstract

In patients with dyssynchronous heart failure (DHF), cardiac conduction abnormalities cause the regional distribution of myocardial work to be non-homogeneous. Cardiac resynchronization therapy (CRT) using an implantable, programmed biventricular pacemaker/defibrillator, can improve the synchrony of contraction between the right and left ventricles in DHF, resulting in reduced morbidity and mortality and increased quality of life. Since regional work depends on wall stress, which cannot be measured in patients, we used computational methods to investigate regional work distributions and their changes after CRT. We used three-dimensional multi-scale patient-specific computational models parameterized by anatomic, functional, hemodynamic, and electrophysiological measurements in eight patients with heart failure and left bundle branch block (LBBB) who received CRT. To increase clinical translatability, we also explored whether streamlined computational methods provide accurate estimates of regional myocardial work.

We found that CRT increased global myocardial work efficiency with significant improvements in non-responders. Reverse ventricular remodeling after CRT was greatest in patients with the highest heterogeneity of regional work at baseline, however the efficacy of CRT was not related to the decrease in overall work heterogeneity or to the reduction in late-activated regions of high myocardial work. Rather, decreases in early-activated regions of myocardium performing negative myocardial work following CRT best explained patient variations in reverse remodeling. These findings were also observed when regional myocardial work was estimated using ventricular pressure as a surrogate for myocardial stress and changes in endocardial surface area as a surrogate for strain. These new findings suggest that CRT promotes reverse ventricular remodeling in human dyssynchronous heart failure by increasing regional myocardial work in early-activated regions of the ventricles, where dyssynchrony is specifically associated with hypoperfusion, late systolic stretch, and altered metabolic activity and that measurement of these changes can be performed using streamlined approaches.

## Introduction

In patients with dyssynchronous heart failure (DHF) characterized by dilated cardiomyopathy (DCM) and electrical conduction delay such as left bundle branch block (LBBB), cardiac resynchronization therapy (CRT) can result in reduced morbidity and mortality and increased quality of life (1) by improving the synchrony of contraction between the right and left ventricles. However, there remains a high non-response rate, and identifying likely CRT responders is difficult (2). A meta-analysis of the major CRT response trials (3) found 71% of recipients were clinical responders as defined by an improvement in symptomatic score, and only 62% were echocardiographic responders as defined by a decrease in left ventricular (LV) end-systolic volume of >10% after 3-6 months, which is indicative of reverse heart failure remodeling (4). Evidence showing a positive relationship between the acute hemodynamic effects of CRT and the long-term clinical and reverse remodeling responses is lacking (5,6). Hence, the mechanisms of short-term and long-term responses to CRT are likely to be different.

The therapeutic mechanism of CRT has been thought to be improved global mechanical pump efficiency due to increased stroke work and decreased myocardial oxygen demand in DHF patients’ contraction (7,8). However, this hypothesis has been questioned with the observation in animal models that CRT reverses many of the spatially heterogeneous molecular and cellular impairments coinciding with redistribution of myocardial workload induced by electromechanical dyssynchrony in LBBB. This redistribution is marked by low or negative work in the early-activated right ventricle (RV) and septum, and elevated work in late-activated LV lateral wall. This results in acute redistribution of myocardial perfusion (9) away from early-activated regions and into late-activated myocardium with corresponding metabolic alterations (10). With chronic dyssynchrony, the wall hypertrophies asymmetrically, thinning and dilating in early-activated regions and thickening in late-active myocardium (11). However, the significance of regional work distributions in human DHF and their changes with CRT are not known, because they depend on the regional myocardial wall stresses, which cannot be directly measured in patients.

To investigate regional work distributions in humans, we used patient-specific, multi-scale computational models (**Fig. 1**) of DHF to compute regional distributions of myocardial work and the changes with biventricular pacing. CRT increased the mechanical efficiency of ventricular pumping, but this improvement was only significant for those patients who showed no evidence of long-term reverse ventricular remodeling at 6-month follow up. In contrast, patients with the greatest heterogeneity of regional myocardial work at baseline showed the greatest reverse remodeling, and the degree of remodeling correlated strongly with the reduction in the fraction of septal myocardium performing negative work. Building patient-specific models is challenging clinically due to the need for numerous different and contemporaneous datasets. Therefore, we tested the accuracy of different approximations to simplify estimation of myocardial work distributions. We found that estimating regional strain as the change in endocardial surface area (12) and regional stress using ventricular pressure led to a regional myocardial work estimate with a strong correlation with model values. However, simplifications which eliminated patient-specific pressure information (use of a generic LV waveform without peak pressure) significantly reduced accuracy.

**Fig. 1:**
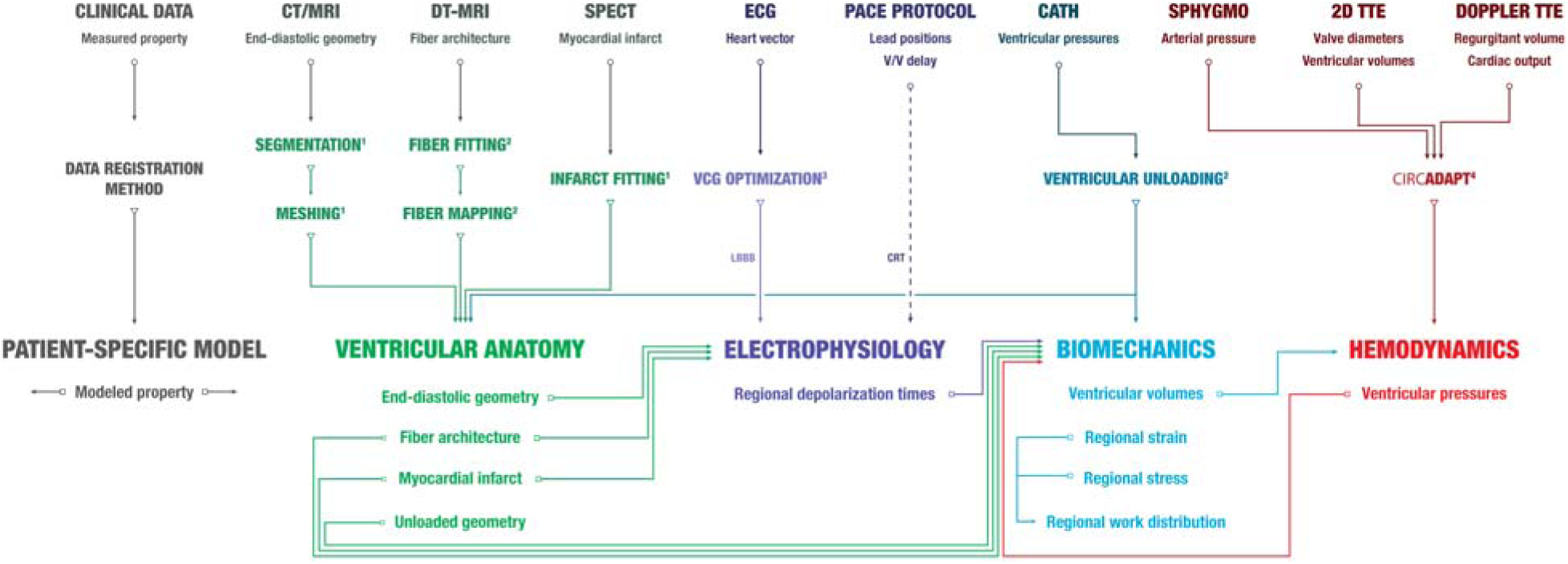
Clinical data, registration methods, and patient-specific model workflow. A patient-specific model includes ventricular anatomy (yellow), electrophysiology (purple), biomechanics (blue) and hemodynamics (red) components. Measured properties from clinical data obtained at baseline were incorporated into corresponding model components using previously described data registration methods (^1^Aguado-Sierra, et al. (18); ^2^Krishnamurthy, et al. (19); ^3^Villongco, et al. (22)). Input/output relationships of model properties between model components are illustrated with arrows. See the online supplement for more details on the patient-specific model.

These new findings suggest that while there is an acute improvement in global ventricular efficiency following CRT, the long-term beneficial effects are related to the normalization of work in early-activated regions having work done on them during systole. These findings suggest that criteria for optimizing CRT pacing based on regional rather than global work alterations may lead to better clinical outcomes. Further, streamlined approaches can be used to accurately estimate regional myocardial work which could improve clinical utility and enable evaluation in larger cohorts.

## Methods

### Clinical Study

Eight male patients (**Table 1**) aged (66±11 years) with NYHA class III heart failure, dilated cardiomyopathy, LBBB (QRS duration 145±22 ms), and left ventricular ejection fraction (LVEF) 28±7% were enrolled from the Veteran’s Administration San Diego Healthcare System (San Diego, CA). Patients gave informed consent to participate in the human subject protocol approved by the Institutional Review Board. Anatomical, electrophysiological, and hemodynamic measurements for each patient were obtained at baseline LBBB and immediately after biventricular cardioverter defibrillator (St. Jude Medical) device implantation and activation (acute CRT). Details can be found in the online supplement (Supplemental Methods). After 6 months of pacing, changes in cardiac output and ventricular volumes (LV reverse remodeling) were assessed by a follow-up 2D transthoracic echocardiography (TTE) study. CRT clinical responders were identified by >1 NYHA class decrease, and echocardiographic responders were identified by >10% decrease in LV end-systolic volume (+ΔESV_LV_) from baseline (4).

**Table 1:** Summary of patient characteristics and measurements at baseline (LBBB) and outcomes (>6 months CRT).

| Measure | BiV1 | BiV2 | BiV3 | BiV4 | BiV5 | BiV6 | BiV7 | BiV8 |
| --- | --- | --- | --- | --- | --- | --- | --- | --- |
| Baseline |  |  |  |  |  |  |  |  |
| Age | 84 | 65 | 79 | 64 | 61 | 55 | 68 | 54 |
| NYHA Class | III | IV | III | II | III | II | IV | III |
| fEF (%) | 38 | 26 | 35 | 25 | 17 | 32 | 21 | 29 |
| QRSd (ms) | 156 | 148 | 162 | 130 | 182 | 119 | 140 | 120 |
| MRV (mL) | — | 16 | — | 36 | 14 | — | — | — |
| HR (bpm) | 70 | 70 | 70 | 70 | 60 | 70 | 70 | 75 |
| Infarct Location | — | septo posterior | septal | posterior | septo posterior | — | — | posterior |
| Outcomes |  |  |  |  |  |  |  |  |
| NYHA Class | II | I | II | I | II | I | IV | II |
| ΔESV<sub>LV</sub> (%) | 12 | -2 | 16 | -12 | 15 | 59 | -36 | 7 |

Patient characteristics include age (years), NYHA classification, forward ejection fraction (fEF) (%), QRS duration (QRSd) (ms), mitral valve regurgitant volume (MRV) (mL), heart rate (HR) (beats per minute), and anatomical location of myocardial infarct. Clinical and echocardiographic responses were determined by the change in the NYHA class and the reduction in LV end-systolic volume (%), respectively.

### Patient Specific Cardiovascular Model

The patient-specific cardiovascular model has four major components: cardiac anatomy, electrophysiology, biomechanics, and hemodynamics. Each component was parameterized from clinical and empirical measurements (**Fig. 1**). See the online supplement (Supplemental Methods) for more details on the model compositions and methods.

Biventricular anatomical finite element models were constructed from a tomographic imaging (CT or MRI) exam (13) with ventricular fiber architecture included by mapping diffusion tensor MR data from a human cadaver heart to each patient model (14). Regions of myocardial infarction were localized from rest and stress SPECT and manually demarcated on the finite element models by an expert cardiac electrophysiologist and nuclear radiologist.

Action potential propagation was modeled using monodomain theory with a human ventricular myocyte ionic current model (15). Patient-specific 3D regional activation times at baseline were estimated by optimizing myocardial electrical conductivities and an ectopic stimulus located in the RV sub-endocardium to match vectorcardiograms derived from measured ECGs as described previously (16). Acute CRT depolarization patterns for paced beats were simulated by applying stimuli in the model using the V-V lead delays and locations selected for each patient.

To simulate ventricular biomechanics, passive material parameters were estimated to match observed end-diastolic pressure and volume measured by intra-cardiac catheterization and echocardiography. Active contractile parameters were optimized to match left and right peak systolic pressure, dP/dt_max_, and dP/dt_min_, and end-systolic volumes as described previously (13). The onset of regional contraction was defined by patient-specific electrical activation times from the electrophysiology model.

Hemodynamics were simulated using a closed-loop, adapting model of the systemic and pulmonary circulation (CircAdapt, (17)) coupled to the boundary conditions of the ventricular model. The fully parameterized hemodynamic model was simulated for 11 beats at measured heart rates until steady-state pressure traces were obtained.

### Global and Regional Myocardial Work Metrics

The external myocardial work density (kJ/m^3^) over the complete cardiac cycle beat was calculated by computing the areas enclosed by stress-strain loops; positive work (counter-clockwise) loops indicate external work performed by the myocardium, and negative (clockwise) work loops define work done on the myocardium. We computed total stroke work, total LV myocardial work, their ratio the efficiency η, the coefficient of variation of work (COVW) and regional RV, septal, and LV work distributions in responders and non-responders during LBBB and CRT.

### Sensitivity Analysis

We performed “knock-out” simulations (6 for LBBB, 2 for CRT) where patient-specific parameters of individual model components were replaced by patient-averaged values across all patients to assess the sensitivity of work metrics to inter-patient variations in physiological properties. The recomputed work metrics from each knock-out set were regressed against ΔESV_LV_ and the original fully-patient-specific model results to assess the relative importance of each property in determining patient variations that contributed to reverse remodeling. More details of the sensitivity analysis are provided in the online supplement (Supplemental Methods).

### Simplified Myocardial Work Estimates

Several simpler approaches to estimate myocardial work were compared to myocardial work calculated with the patient-specific model to assess agreement with and the need for advanced patient-specific modeling. Estimates were calculated as the areas enclosed by approximations of stress-strain loops. For all estimates, strain was approximated by triangulating the endocardial points on the patient-specific geometric model and tracking changes to the area of each triangle throughout the cardiac cycle (12,18). For a patch v, strain was calculated as:

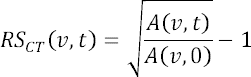

where *t* is the temporal position in the RR interval and *A* is the area of patch *v* (12). This approach has been successfully used to characterize regional endocardial strain using ECG-gated cine CT images in patients with normal function (19), with abnormal function (20,21), and undergoing interventional therapy (22). To assess segmental strain, *RS_CT_* was averaged within AHA segments to achieve one strain curve per segment. Stress was approximated using the methods detailed below.

Prior studies have utilized the ventricular pressure-strain loop area (PSA) (10, 23–25), with LV pressure acquired from left heart catheterization, as a surrogate for regional myocardial work. The LV pressure provides some patient-specific information (timing and peak value) but eliminates regional variation in stress, as the same LV pressure waveform was applied to all strain curves. This estimate is referred to as the P_LHC_SA in the results.

As described above, ventricular pressure as a surrogate for myocardial stress overlooks regional heterogeneities which may influence regional myocardial work. Therefore, we tested two geometric approaches to incorporate local shape information. The second and third myocardial work estimates were measured as the end-diastolic wall stress-strain loop area (WS_ED_SA) and time-varying wall stress-strain loop area (WS_TV_SA), respectively. Wall stress, σ, was calculated with the Laplace law (26):

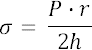

where *P* is the LV pressure from catheterization, *r* is the segmental radius, and *h* is the segmental wall thickness. Segmental LV wall thickness was derived from each patient-specific model. Segmental radius was calculated by fitting an ellipsoid to the endocardial points of the geometric patient-specific model that represent each LV segment. To account for circumferential and longitudinal radius, segmental radius was calculated as the effective radius:

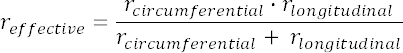

surface curvature K can be described as a function of k_longitudinal_ + k_circumferential_ (27). Given that radius is the inverse of curvature, converting K ≍ k_longitudinal_ + k_circumferential_ in terms of radius yields r_effective_.

The static wall stress for WS_ED_SA was calculated with principal radius and wall thickness at end-diastole. This creates a constant scaling factor for each AHA segment which is then applied to the patient’s LV pressure waveform. The time-varying wall stress for WS_TV_SA method leads to a scaling factor which changes during the cardiac cycle.

Finally, we evaluated the need for patient-specific pressure recordings. Two estimates were generated by using a generic pressure waveform digitized from a Wigger’s diagram of normal hemodynamics (Human Bio Media). The fourth estimate applied the generic normal LV pressure waveform with each strain curve (P_gen_SA). In the fifth estimate, we scaled the generic waveform scaled such that the peak pressure achieved the same patient-specific peak pressure acquired with catheterization (P_gen,scaled_SA).

All myocardial work estimates are reported in terms of kPa; all stress estimates are in kPa and the strain estimate is unitless. Agreement between segmental strain and myocardial work measured with limited clinical data and patient-specific modeling is described in **Supplement: Simplified Work Estimates**. To evaluate the utility of these simplified myocardial work estimates in predicting CRT response, we computed the fraction of negative work relative to the endocardial surface area in the LV (S_f_LVNW) and septum (S_f_STNW) during LBBB in known CRT responders and non-responders.

### Statistics

Average data are reported as mean ± SD. Significant (p<0.05) differences in work metrics were determined by student’s t-tests and 2-way ANOVA between responders versus non-responders and LBBB versus CRT. Work metrics that differed significantly between response groups and physiologic states were correlated with LV reverse remodeling by simple linear regression with significance computed by the critical values for Pearson’s correlation coefficient.

## Results

### Clinical Measurements Do Not Correlate with Clinical Outcomes

Clinical outcomes of the eight patients implanted with biventricular pacemaker-defibrillators were assessed after approximately six months of pacing (7.3±4.7 months). Four were echocardiographic CRT responders as defined by a reduction in LV end-systolic volume (ΔESV_LV_) > 10% (**Table 1**); ESV_LV_ in the best responder (BiV6) decreased by more than 50%, whereas in three of the four echocardiographic non-responders ESV_LV_ increased after therapy. One of these three (BiV7) was classified as a clinical non-responder, while the other seven all showed improvement of ≥ 1 NYHA heart failure class.

None of the baseline clinical measurements including QRS duration, LV ejection fraction (LVEF), LV end-diastolic volume, LV peak systolic pressure, dP/dt_max_, dP/dt_min_, LV end-diastolic volume, and LV mass correlated with ΔESV_LV_ with an R^2^ > 0.2 (**Fig. 2**).

**Fig. 2:**
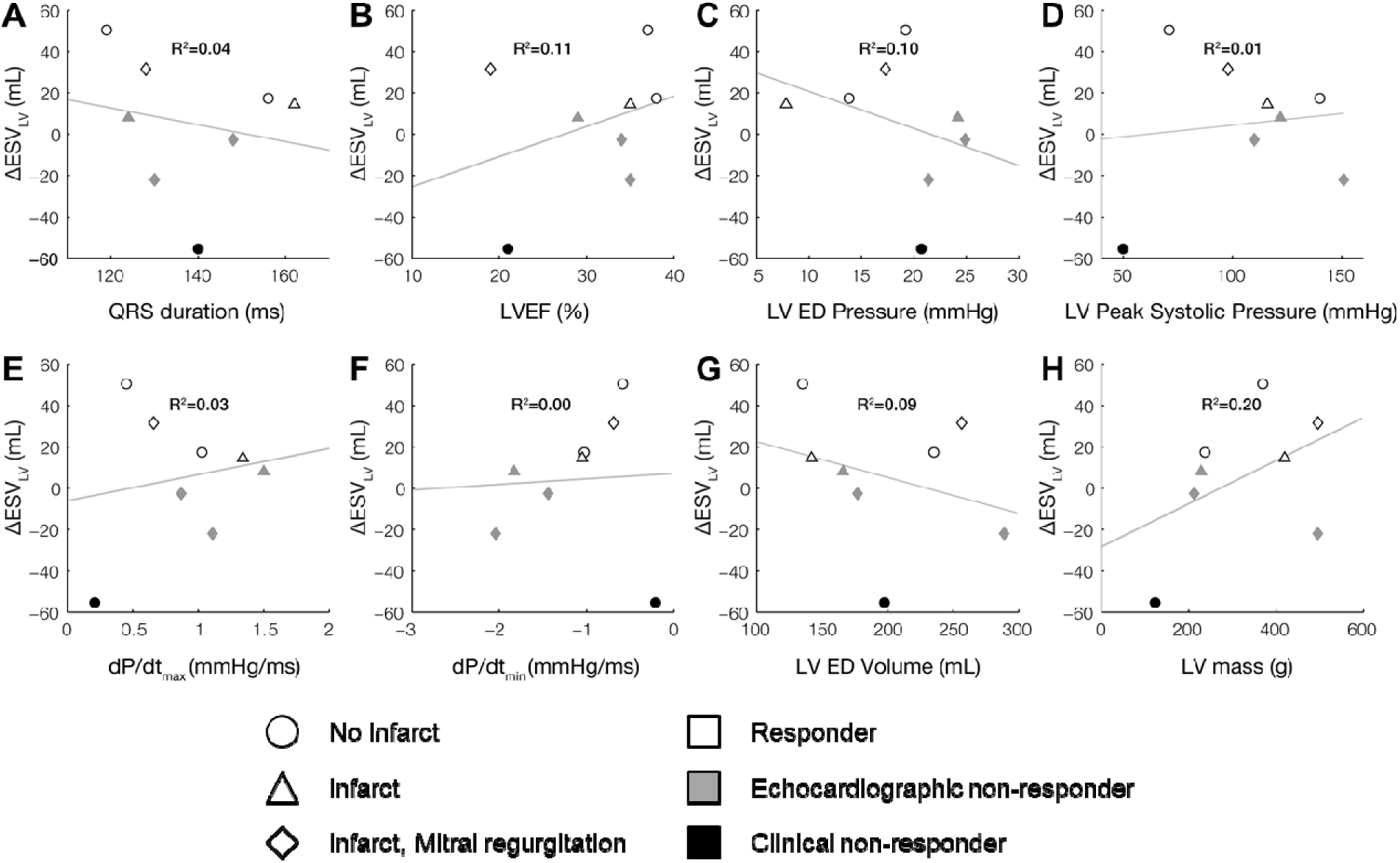
Clinical measures of LV function at baseline (LBBB) do not correlate with CRT outcome. Baseline clinical measurements including (A) QRS duration, (B) left ventricular ejection fraction (LVEF), (C) LV end-diastolic (ED) pressure, (D) LV peak systolic pressure, (E) dP/dt_max_, (F) dP/dt_min_, (G) LV ED volume, and (H) LV mass do not correlate with measured ΔESV_LV_ after 6 months of CRT.

### Acute Hemodynamic and Electrophysiological Alterations after CRT

When CRT pacing was applied to the baseline patient-specific models, the computed changes in peak LV systolic pressure and dP/dt_max_ agreed well with the values measured acutely after the pacemaker was switched on at the time of the implantation procedure (**Fig. 3**). RMS errors between the model-predicted and measured peak LV pressure and dP/dt_max_ were 5 mmHg and 0.3 mmHg/s, respectively. Model-computed QRS durations during CRT pacing differed from the values measured in patients after the pacemaker was turned on by an RMS error of 30 ms.

**Fig. 3:**
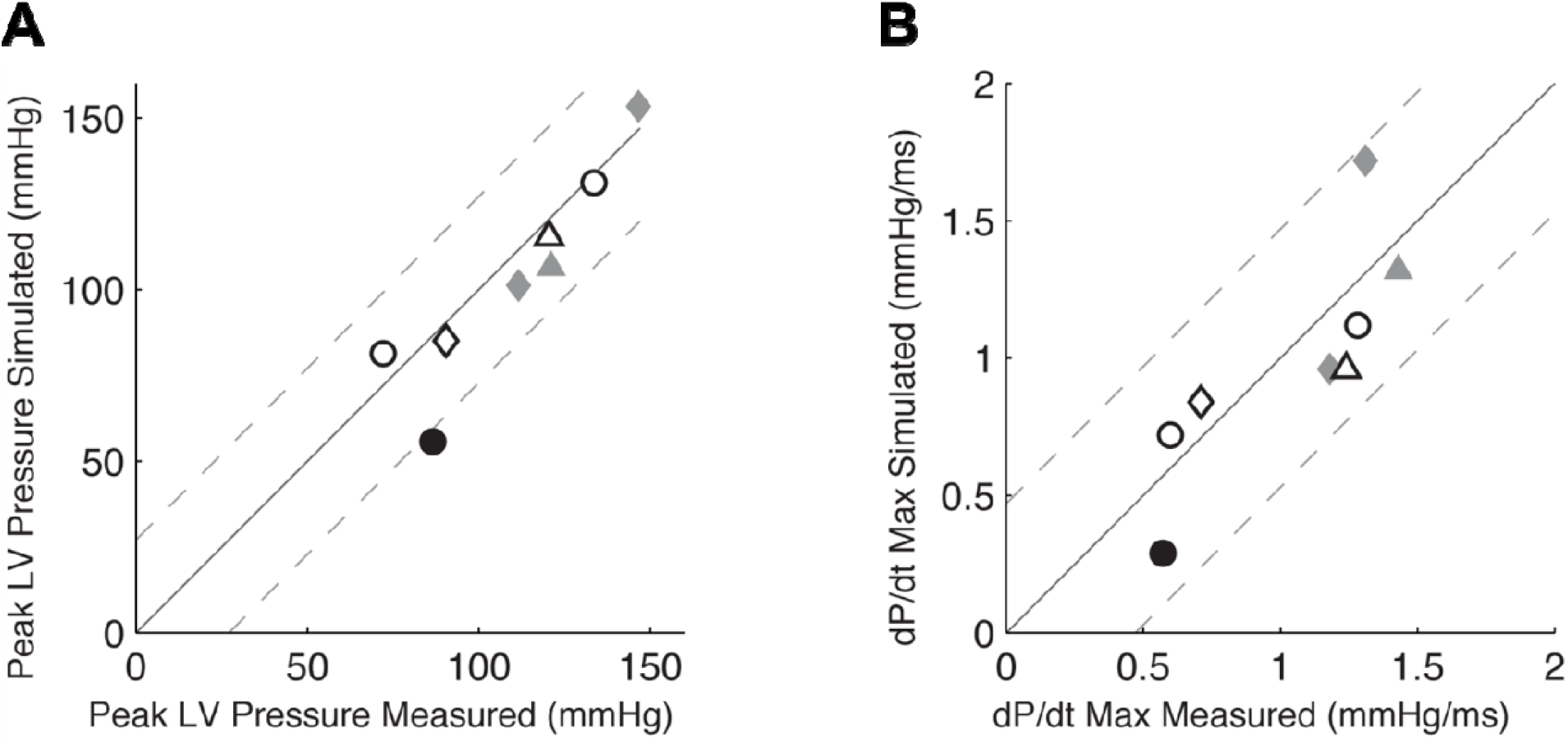
Patient-specific CRT model validation. Model-predicted dP/dt_max_ (A) and peak systolic pressure (B) during CRT compared with measured values in patients acutely after pacing initiation.

### CRT Improved Ventricular Pumping Efficiency Computed with Patient-Specific Models

After CRT, model-computed LV stroke work (**Fig. 4A**) increased (by 0.05±0.04 J in responders and 0.01±0.003 J in non-responders) and model-computed total myocardial work (**Fig. 4 AB)** decreased (by 0.002±0.05 J in responders and 0.05±0.01 J in non-responders). These small acute changes following CRT were not statistically significant, however ventricular mechanical efficiency η—the ratio of global LV stroke work to total LV myocardial work— increased by 7.7±1.4% (**Fig. 4C**), and this difference was statistically significant (p<0.005), but only in the echocardiographic non-responders (8.6±2.0%). Therefore, while CRT can increase global ventricular mechanical energy efficiency, this acute improvement does not appear to explain the longer-term beneficial effects of resynchronization.

**Fig. 4:**
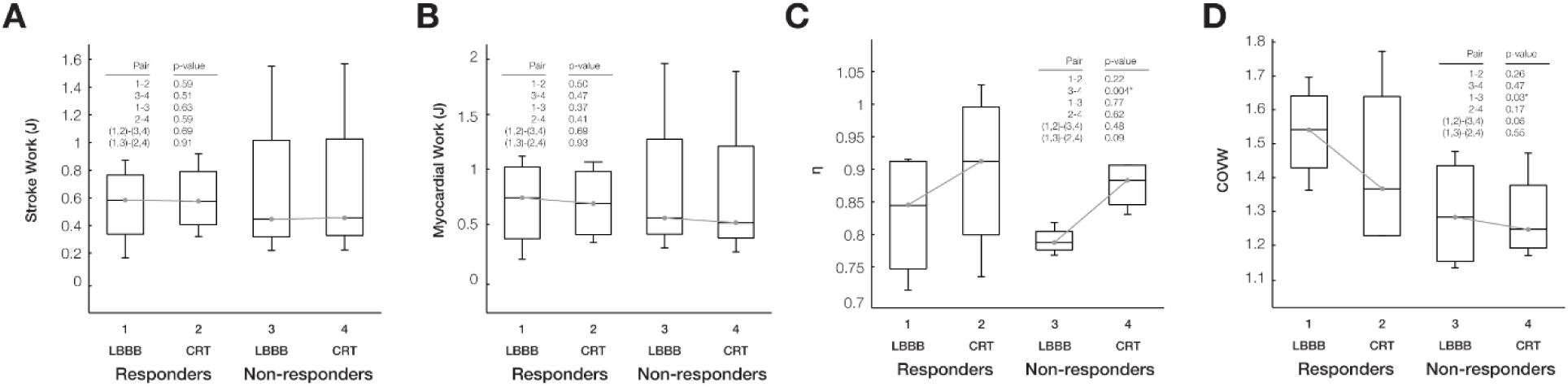
Regional work heterogeneity in LBBB differs between responders and non-responders, but LV stroke work, myocardial work, and efficiency do not. Box plots of computed distributions of (A) stroke work (J), (B) myocardial work (J), (C) mechanical efficiency (η), and (D) coefficient of variation of work (COVW). Changes in stroke work, and total myocardial work after CRT were not significant, whereas η increased after CRT though this change was only significant for non-responders. (D) Regional work heterogeneity (COVW) was significantly (p<0.05) higher at baseline in responders than non-responders and only decreased significantly after CRT (p=0.05) in the responder group.

### Baseline Regional Work Heterogeneity was Greater in Responders than Non-Responders

The coefficient of variation of regional myocardial work density (COVW) computed from regional fiber stress-strain loops throughout each patient-specific model (**Fig. 4D**) was significantly higher in responders (1.53±0.07) than non-responders (1.29±0.08) at baseline (p<0.05). The decrease in work heterogeneity due to simulated CRT was greater in responders (0.10±0.09) than in non-responders (0.01±0.04), but the difference between the groups was not statistically significant.

Differences in COVW between responders and non-responders were significant when derived from simplified estimates of regional myocardial work via ventricular pressure (P_LHC_SA: 0.99 ± 0.17 and 0.61 ± 0.17, p = 0.02), a generic pressure waveform (P_gen_SA: 1.26 ± 0.37 and 0.71 ± 0.11, p = 0.03), and a scaled waveform (P_gen,scaled_SA: 1.26 ± 0.37 and 0.71 ± 0.11, p = 0.02) (**Fig. 5**). However, shape-informed work estimates WS_ED_SA (1.12 ± 0.16 and 1.25 ± 0.87, respectively, p = 0.77) and WS_TV_SA (1.19 ± 0.21 and 1.13 ± 0.56, respectively, p = 0.86) did not separate responders and non-responders.

**Figure 5:**
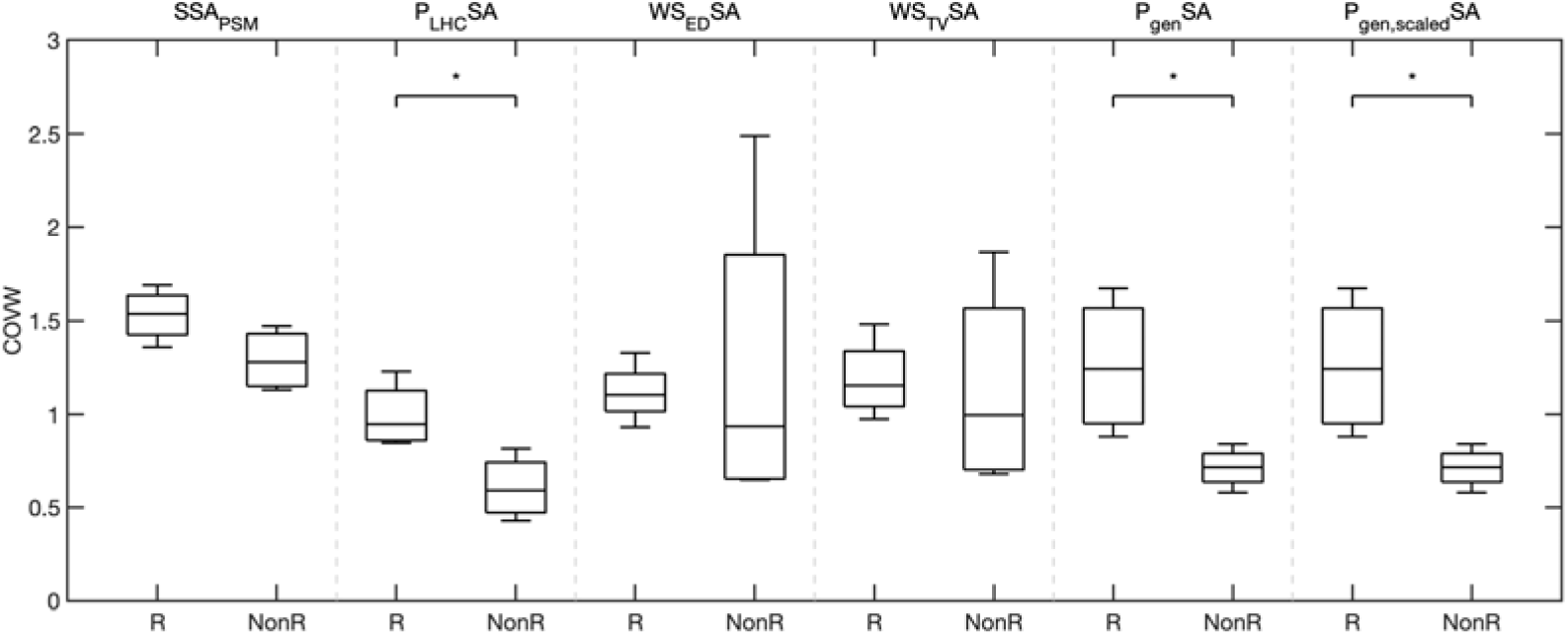
Regional work heterogeneity in LBBB differs between CRT responders and non-responders for work estimated as the pressure-strain loop. Box plots of the coefficient of variation of work (COVW) computed with different myocardial work estimates: stress-strain area from patient-specific modeling (SSA_PSM_), catheter derived LV pressure-strain area (P_LHC_SA), end-diastolic Laplace wall stress-strain area (WS_ED_SA), time-varying Laplace wall stress-strain area (WS_TV_SA), generic LV pressure-strain area (P_gen_SA), and peak pressure-scaled generic LV pressure-strain area (P_gen,scaled_SA). COVW during LBBB is significantly higher in responders than non-responders when computed with pressure-strain area approaches (P_LHC_SA, P_gen_SA, P_gen,scaled_SA), but not when COVW is computed with shaped-informed approaches (WS_ED_SA and WS_TV_SA). *indicates significance (p<0.05)

### Regional Work Distributions Differ Between Responders and Non-responders

Patient-specific model-computed regional work distributions showed greater heterogeneity at baseline and greater uniformity after CRT in the best responder than the worst non-responder (**Fig. 6**). Regional fiber stress-strain loops show regions of low or negative loop area (where loops cross in a figure-8 morphology during systole) in early-activated regions (the RV free wall and septum) prominently in the responder, but not in the non-responder. In contrast, late-activated regions in the LV free wall of both patients produced large loops with abnormally high positive work magnitudes due to increased systolic stress. This region of high regional work was reduced more in the responder than in the non-responder.

**Fig. 6:**
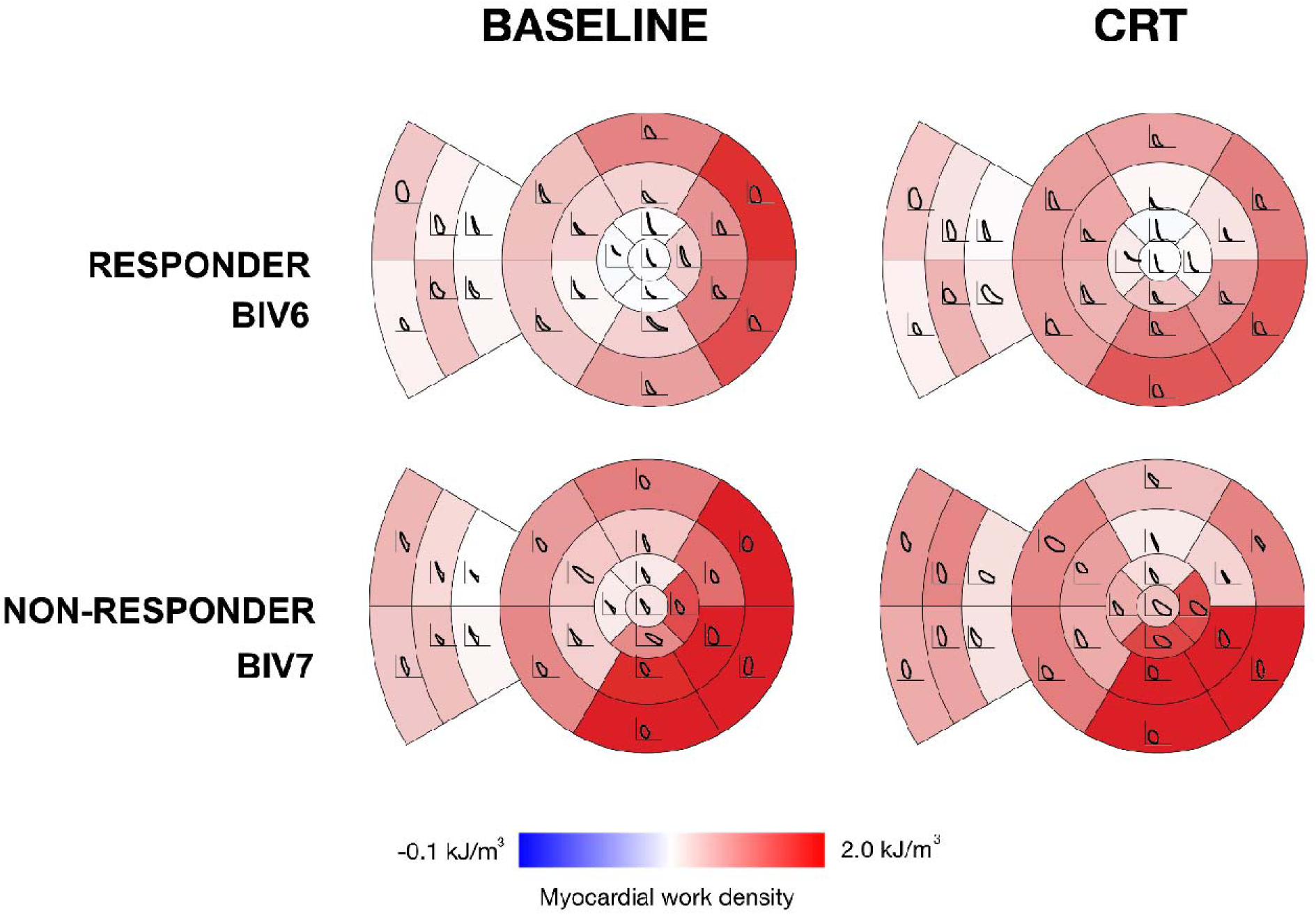
Myocardial regional work density distributions in one responder and one non-responder before and after CRT. Myocardial work density was computed from simulated fiber stress-strain loops during LBBB and CRT for the best responder (top) and worst non-responder (bottom). The LV myocardial work density shown in color for standard myocardial segment with fiber stress-strain loops overlaid. Note that model-computed regional work is more heterogeneous in the responder than in the non-responder at baseline and less heterogeneous in the responder than in the non-responder acutely after CRT.

Myocardial volume distributions of regional work density in the LV free wall, septum, and RV free wall regions at baseline (**Fig. 7A**) and after CRT (**Fig. 7B**) showed long tails of high work regions, primarily in the LV free wall, and short tails of negative work regions, particularly in the RV free wall and septum. At baseline, the volume fraction of the left ventricle performing high regional work (upper quartiles of work density distributions) was not significantly different between responders and non-responders (0.30±0.01 vs. 0.31±0.02). CRT reduced the volume fraction of the left ventricle performing high regional work, but the magnitude of this reduction was not significant in either echocardiographic non-responders (0.040±0.005, p=0.12) or responders (0.005±0.004, p=0.34).

**Fig. 7:**
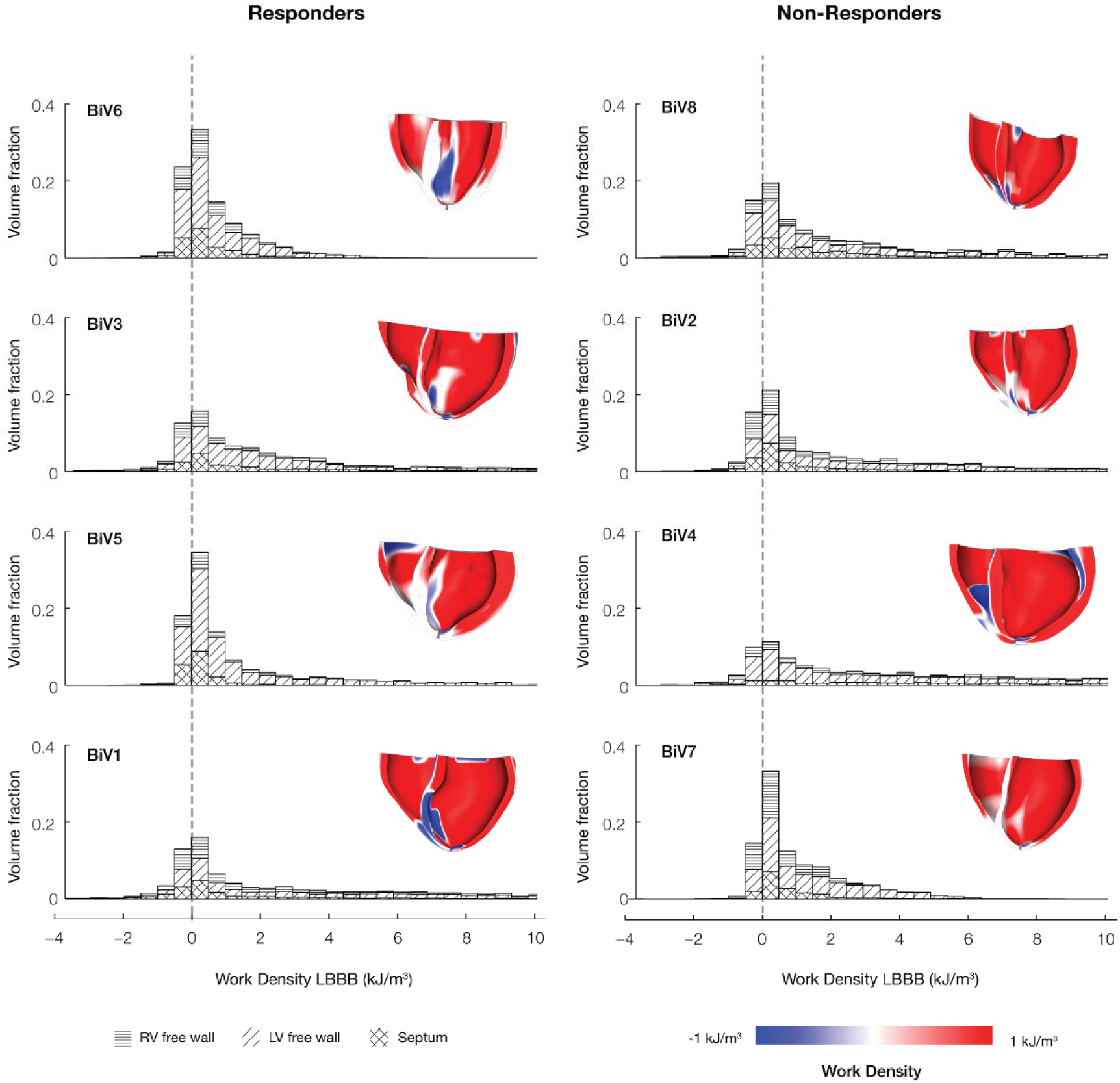
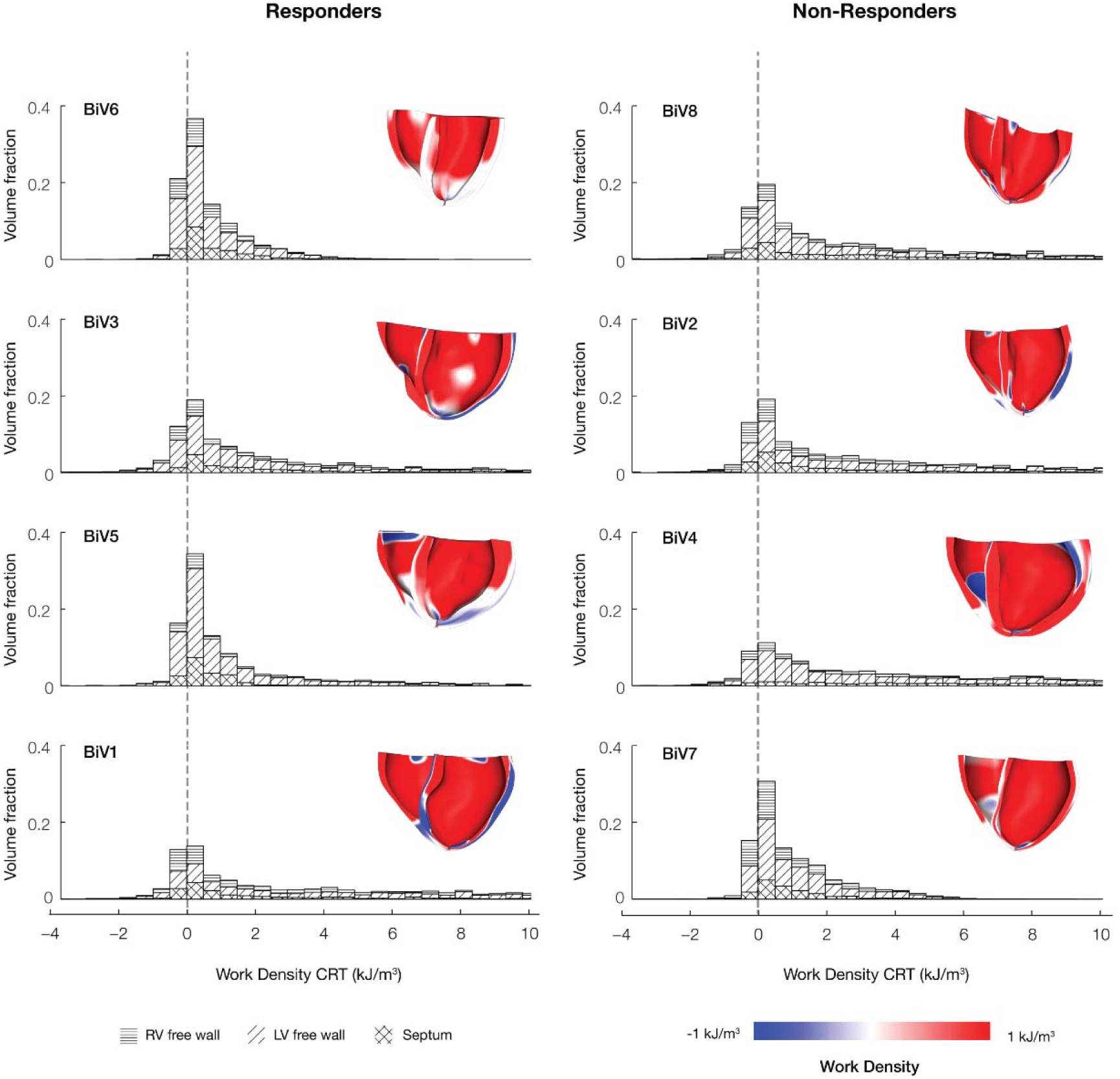
Regions of the LV and septum perform negative work in LBBB and CRT. The regional myocardial volume distributions myocardial work density segregated by septum (cross-hatched), LV free wall (diagonal hatched), and RV free wall (horizontal hatched) regions. The fraction of myocardium performing negative work was associated with the degree of measured CRT reverse remodeling response (ΔESV_LV_). Patients are ordered column-wise according to degree of measured improvement (BiV6 was the strongest responder, and BiV7 was the weakest responder).

The volume fractions of left ventricle and septum performing negative work (V_f_LVNW and V_f_STNW, respectively) were significantly higher at baseline in responders (0.19±0.04 and 0.28±0.03, respectively) than in non-responders (0.14±0.03 and 0.17±0.03, respectively, **Fig. 8A** and **8B**). The reduction in V_f_LVNW and V_f_STNW after CRT was significant in responders (0.19±0.04 to 0.15±0.01, p<0.05 and 0.28±0.03 to 0.12±0.02, p<0.01, respectively) but not in non-responders. The differences in the mean changes in these quantities after CRT between responders (-0.04±02 and -0.16±0.01, respectively) and non-responders (-0.004±0.008 and - 0.05±0.02, respectively) were also significant (p<0.05).

**Fig. 8:**
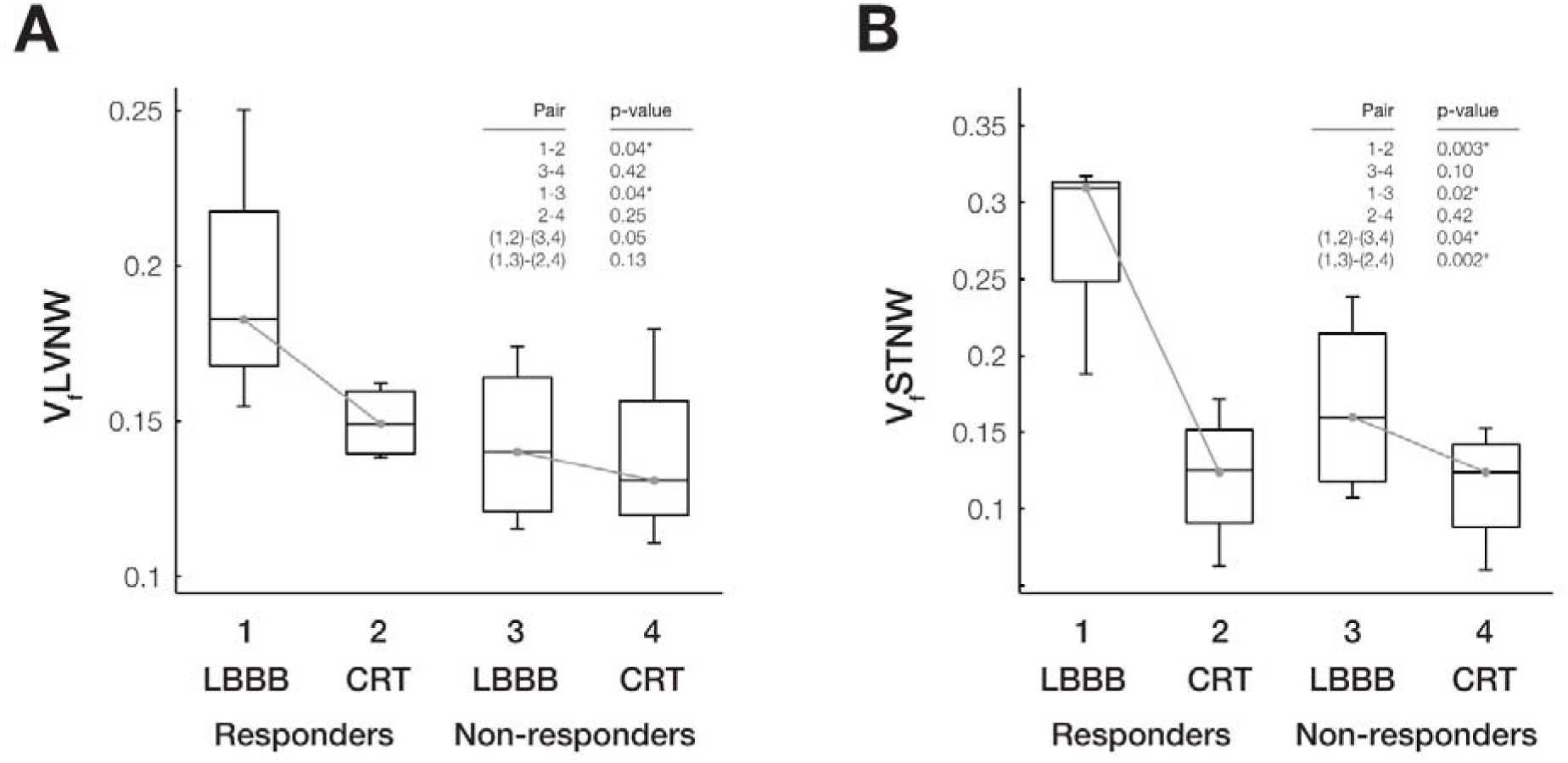
Regional fractions of LV myocardium and septum with negative work differ between responders and non-responders during LBBB and CRT. Box plots of computed distributions of work and metrics show significant (p<0.05) differences between responders and non-responders during LBBB and CRT. The volume fractions of LV with (A) negative work (V_f_LVNW) and (B) septum with negative work (V_f_STNW) are significantly different among responders during LBBB compared with CRT (Pair 1-2) and during LBBB between responders and non-responders (Pair 1-3). STNW is also significantly different for all patients during LBBB and CRT (Pair (1,2)-(3,4) and Pair (1,3)-(2,4)).

Simplified myocardial work measurements estimated S_f_LVNW to be significantly higher in responders (P_LHC_SA: 0.18 ± 0.09; WS_ED_SA: 0.18 ± 0.09; WS_TV_SA: 0.18 ± 0.09; P_gen_SA: 0.25 ± 0.12; P_gen,scaled_SA: 0.25 ± 0.12) than non-responders (P_LHC_SA: 0.06 ± 0.05; WS_ED_SA: 0.08 ± 0.08; WS_TV_SA: 0.07 ± 0.08; P_gen_SA: 0.09 ± 0.04; P_gen,scaled_SA: 0.09 ± 0.04) (**Fig. 9**).

**Figure 9:**
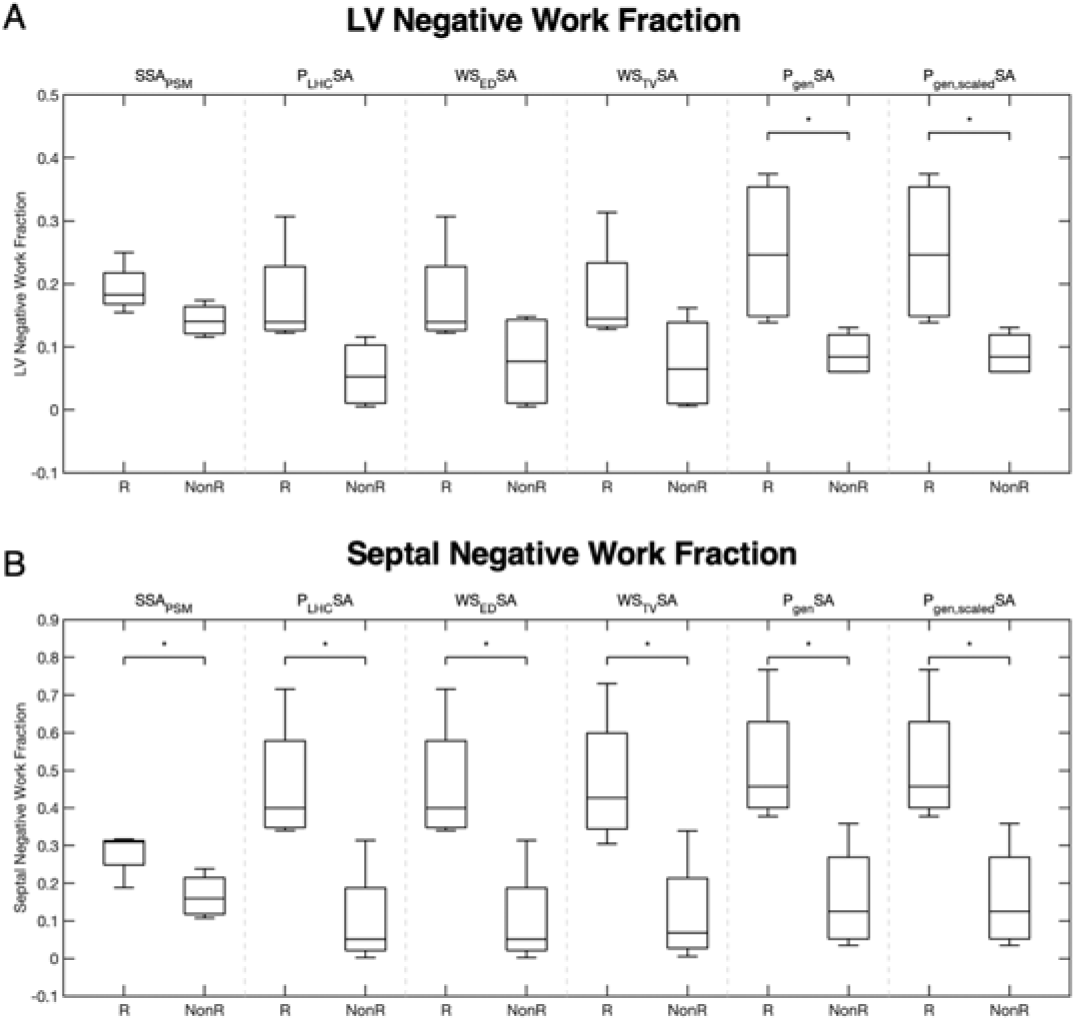
Regional fractions of LV and septal endocardial negative work measured with simplified estimates differ between CRT responders and non-responders. Box plots of the extent of negative work in the LV (A) and septum (B) computed with different myocardial work estimations: stress-strain area from patient-specific modeling (SSA_PSM_), catheter derived LV pressure-strain area (P_LHC_SA), end-diastolic Laplace wall stress-strain area (WS_ED_SA), time-varying Laplace wall stress-strain area (WS_TV_SA), generic LV pressure-strain area (P_gen_SA), and peak pressure-scaled generic LV pressure-strain area (P_gen,scaled_SA). Regional fractions of the LV and septum performing negative work in LBBB are significantly higher in responders than non-responders for all work estimates. Note that negative work found with SSA_PSM_ was computed as the volume fraction of LV and septum with negative work (V_f_LVNW and V_f_STNW), while the simplified work measurements computed the surface fraction of negative work on the LV (S_f_LVNW) and septum (S_f_STNW). Label “R” denotes CRT responders and “NonR” denotes non-responders. *indicates significance (p< 0.05).

Simplified myocardial work measurements also estimated S_f_STNW to be higher in responders (P_LHC_SA: 0.46 ± 0.17; WS_ED_SA: 0.46 ± 0.17; WS_TV_SA: 0.47 ± 0.18; P_gen_SA: 0.51 ± 0.17; P_gen,scaled_SA: 0.51 ± 0.17) than non-responders (P_LHC_SA: 0.10 ± 0.14; WS_ED_SA: 0.10 ± 0.14; WS_TV_SA: 0.12 ± 0.15; P_gen_SA: 0.16 ± 0.15; P_gen,scaled_SA: 0.16 ± 0.15) (**Fig 9**).

### Reduction in Volume Fraction of the Septum Performing Negative Work after CRT Predicts LV Reverse Remodeling

The volume fractions of the left ventricular and septal regions performing negative myocardial work at baseline computed with the patient-specific models correlated strongly and significantly with the measured ΔESV_LV_ after 6 months (**Fig. 10 A and B**). At baseline, V_f_LVNW explained 82% of the variation in ΔESV_LV_ between patients (p<0.01) and V_f_STNW explained 68% of the variation in ΔESV_LV_ between patients (p<0.02).

**Fig. 10:**
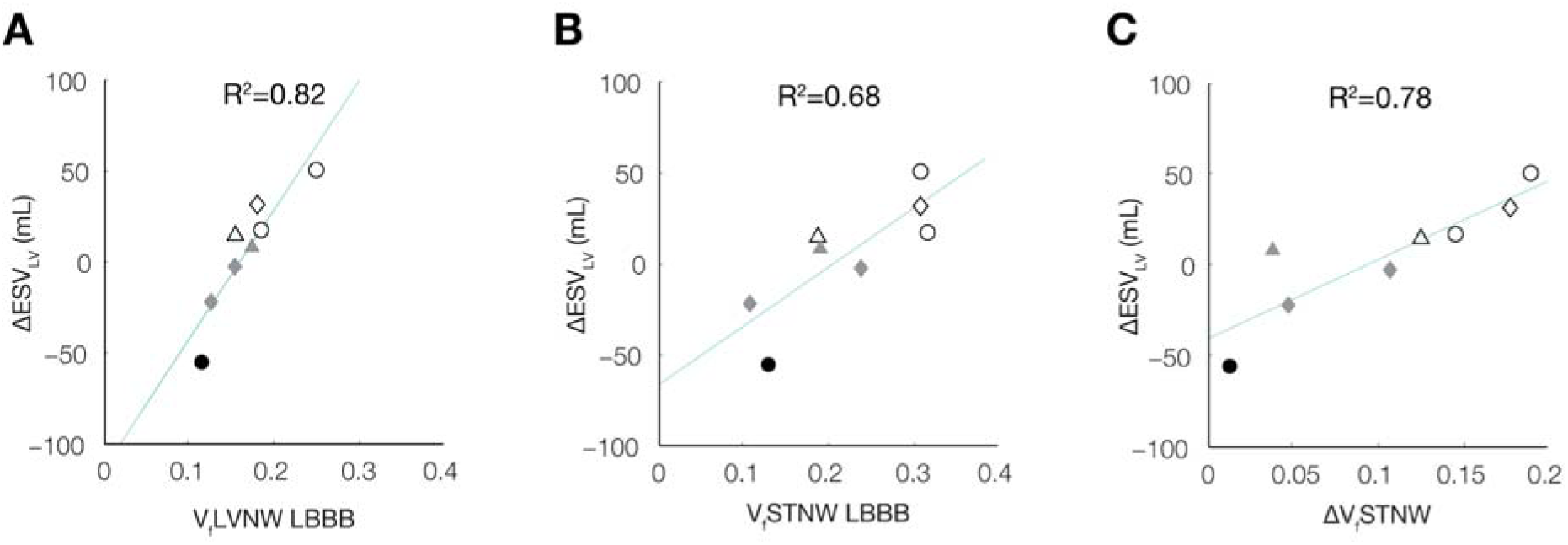
Reduction of negative work in the LV and septum relates to the degree of LV reverse remodeling. Correlations between LV reverse remodeling (ΔESVLV) and volume fractions of the (A) LV with negative work (VfLVNW) during LBBB, (B) septum with negative work (VfSTNW) during LBBB, and (C) ΔVfSTNW (defined as VfSTNWLBBB–VfSTNWCRT) are shown. Shading color denotes patient echo responders (ΔESVLV >10%) (white), echo non-responders (ΔESVLV <10%) (grey), and clinical non-responders (black); shapes denote non-ischemic (circle), ischemic (triangle), and ischemic with mitral regurgitation (diamond). At baseline, larger VfLVNW (A) and VfSTNW (B) are associated with better ΔESVLV outcomes. Moreover, ΔVfSTNW (C) correlates with of ΔESVLV, suggesting improved septal work as an important feature leading to reverse remodeling.

The reductions in the volume fractions of the regions performing negative myocardial work after CRT also correlated significantly with ΔESV_LV_, both for the left ventricle (ΔV_f_LVNW: R^2^=0.52; p<0.05) and the septum (ΔV_f_STNW: R^2^=0.78; p<0.01). Since these regional alterations are predictions of mechanistic models derived from clinical measurements that were not independently correlated with clinical outcomes, these correlations suggest a causal relationship between the reduction in the volume of septal regions performing negative work induced by CRT and the subsequent extent of reverse ventricular remodeling.

The correlations between baseline S_f_LVNW and ΔESV_LV_ after 6 months when regional work was estimated using P_LHC_SA and WS_ED_SA were significant and moderate (R^2^ = 0.51, p < 0.05; R^2^ = 0.50, p < 0.05, respectively). However, WS_TV_SA and estimates using generic waveforms (P_gen_SA and P_gen,scaled_SA) did not significantly correlate. Further, simplified estimates of baseline S_f_STNW did not correlate significantly with ΔESV_LV_ after 6 months. Myocardial work estimated with simplified approaches was not evaluated after CRT, therefore correlations between the change in regional areas performing negative work after CRT and ΔESV_LV_ were not available for the simplified measurements.

### Negative Work Fraction Is Sensitive to Hemodynamics and the Interaction between Ventricular Dilation and Electrical Dyssynchrony

To identify those properties that were most important in determining differences between patients in regional negative work volume fractions and their changes after CRT, patient-specific model parameters were individually replaced by patient-averaged parameters for: baseline ventricular geometry (-G), myocardial infarction (-I), baseline ventricular electrical activation sequence (-A), baseline myocardial resting and contractile material properties (-M), baseline circulatory hemodynamics (-H). In addition to examining the effects of each substitution separately, we examined the combined effects of eliminating patient-specific geometry and activation sequence in the models (-GA). To examine the importance of patient-specific CRT pacing parameters on work distributions, we re-ran CRT models for each patient using the average lead locations and V-V delays. In total, these model perturbations applied individually to each patient resulted in six permutations of each of the original 8 baseline patient-specific models and three permutations of the original 8 CRT simulations (**Table 2**).

**Table 2:** Summary results of work sensitivity analysis.

| State | LBBB |  |  |  |  |  |  | ΔCRT |  |  |
| --- | --- | --- | --- | --- | --- | --- | --- | --- | --- | --- |
| Case | Ps | −G | −I | −A | −M | −H | −GA | Ps | −G | −A |
| Model Component |  |  |  |  |  |  |  |  |  |  |
| Geometry | ● | ○ | ● | ● | ● | ● | ○ | ● | ○ | ● |
| Infarct | ● | ● | ○ | ● | ● | ● | ● | ● | ● | ● |
| Activation times (LBBB) | ● | ● | ● | ○ | ● | ● | ○ | – | – | – |
| Biochanics | ● | ● | ● | ● | ○ | ● | ● | ● | ● | ● |
| Hemodynamics | ● | ● | ● | ● | ● | ○ | ● | ● | ● | ● |
| Activation times (CRT) | – | – | – | – | – | – | – | ● | ● | ○ |
| Regional work metric |  |  |  |  |  |  |  |  |  |  |
|  | R<sup>2</sup> (ΔESV<sub>LV</sub>) |  |  |  |  |  |  |  |  |  |
| V<sub>f</sub>LVNW | 0.815* | 0.580* | 0.823* | 0.733* | 0.797* | **0.374** | **0.478** | 0.516* | **0.087** | **0.183** |
| V<sub>f</sub>STNW | 0.675* | 0.546* | 0.752* | 0.605* | 0.629* | **0.130** | **0.466** | 0.779* | **0.499** | **0.044** |
|  | nRMSD (Ps) |  |  |  |  |  |  |  |  |  |
| V<sub>f</sub>LVNW | 0% | **14%** | 2% | 9% | 3% | **31%** | **14%** | 0% | **69%** | **34%** |
| V<sub>f</sub>STNW | 0% | **33%** | 11% | 13% | 3% | **33%** | **31%** | 0% | **41%** | **47%** |

The sensitivity of computed volume fractions of negative work in the LV ( $V_{fLVNW}$ ) and septum ( $V_{fSTNW}$ ) at baseline and change after CRT to model parameters was assessed by substituting the original patient-specific (**Control**) parameters with patient-averaged parameters for the geometry (**G**), activation times (**A**), mechanics (**M**), and hemodynamics (**H**). In patients with myocardial infarction, the infarct was removed (**I**). Patient-averaged hemodynamics and simultaneous geometry and electrical activation times (**GA**) eliminated the significant correlation between the outcome and the volume fraction of the LV performing negative work at LBBB. The significant correlation between $\Delta ESV_{LV}$ and $\Delta V_{fSTNW}$ was also eliminated using average geometry and electrical activation pattern during acute CRT.

Recomputing the regional distributions of myocardial work density and metrics of work heterogeneity for all 72 cases, we found that patient variations in baseline negative work fractions were only significantly altered in models that excluded patient-specific hemodynamics (-H), or those that excluded both patient-specific geometry and activation sequence (-GA), though not either property alone. Patient-to-patient differences in the reduction in negative myocardial work after CRT were also significantly altered when patient-specific geometry and pacing protocol were eliminated. This analysis suggests that differences between patients in regional negative work fractions and their changes after CRT are most sensitive to patient variations in hemodynamics, ventricular geometry and activation sequence, whereas differences in myocardial systolic and diastolic material properties, the presence of a myocardial infarct or the specific CRT pacing protocol were not primary determinants of CRT outcomes.

## Discussion

This was the first study to investigate distributions and changes of regional myocardial mechanical work in human DHF before and after CRT. This was only possible with the use of multi-scale patient-specific computational models parameterized with comprehensive structural, hemodynamic, electrophysiological, and physiological measurements in patients who were candidates for CRT. Although, as reported previously (28), CRT improved the mechanical efficiency with which regional myocardial work was converted to LV stroke work, there was no relationship between the magnitude of this improvement and the extent of long-term reverse ventricular remodeling. Indeed acutely, efficiency increased significantly more in echocardiographic non-responders than responders. This may help explain the poor correlation between improvements in symptomatic scores post-CRT compared with long-term ventricular reverse remodeling (5). Other heart failure treatments that chronically improve ventricular pump performance and survival typically do not confer acute inotropic benefits, and the current findings suggest that CRT might not be an exception to this observation.

### Experimental evidence of physiological responses to altered workload

Animal studies showing that CRT reverses metabolic and cellular remodeling caused by dyssynchrony have called into question the importance of improved global energy efficiency for CRT response (1). Many of the energetic, metabolic, and stress signaling responses to dyssynchrony are spatially heterogeneous in the ventricular walls. Large animal studies have elucidated the origins of this spatial heterogeneity. Prinzen, et al. (29) showed that RV apical pacing, which models LBBB, results in low or negative regional myocardial work in early-activated regions of the septum and elevated regional work in late-activated regions on the LV free wall. This is associated with an acute redistribution of myocardial perfusion away from the RV and septum and into the LV free wall (30). Reduction of coronary flow reserve has been observed in the septum of non-ischemic LBBB patients (31). More recent studies in DHF patients have demonstrated that depressed septal glucose metabolism at baseline and improvement in septal perfusion during CRT may predicts long-term LV reverse remodeling (32). Nemutlu, et al. (33) recently reported that CRT significantly altered plasma metabolomic profiles consistent with improved substrate oxidation and energy metabolism. These studies suggest that altered myocardial mechanoenergetics may underlie metabolic remodeling associated with DHF pathophysiology and its reversal by CRT. Therefore, we examined regional distributions of myocardial work density before and after CRT using patient-specific models and focused specifically on regional work heterogeneity.

Many animal studies have also shown that dyssynchrony leads to heterogeneous cellular and molecular changes in the left ventricular free wall and septum (9,33,34). Transcripts for proteins involved in metabolic networks (oxidative phosphorylation, fatty acid, amino acid, and glucose metabolism) were down-regulated, primarily in cardiomyocytes sampled from early-activated regions, while components of signaling pathways (MAPK, JAK-STAT, TGF-beta) were up-regulated in late-activated sites (35). CRT has been seen to reverse these heterogeneities including rebalancing of glucose metabolism and restoring myocardial blood flow (36). Here, we found that reverse ventricular remodeling was not predicted by the reduction after CRT in regions of the left ventricular free wall performing high work, but did correlate strongly with the reduction in the size of the septal region performing negative work. This suggests that further studies focusing on the regional alterations in perfusion, metabolism, and cell signaling that occur primarily in these early-activated regions of the myocardium may elucidate important mediators of long-term reverse ventricular remodeling.

### Work heterogeneity is an important property of the total physiological impact of electromechanical dyssynchrony

At baseline, regional work was highly heterogeneous in patients with LBBB, as measured by the COVW. In addition, those patients with greatest work heterogeneity tended to be the best responders to CRT. These findings confirm in humans the conclusions of Kerckhoffs, et al. (37), who modeled canine DHF, that COVW is a sensitive measure of the severity of dyssynchrony in the failing heart. Examining these distributions in more detail, the long-term response to CRT was not, however, explained by the acute reduction in overall work heterogeneity *per se,* or in the decrease in the size of the late-activated regions performing high regional work. The models did show that the reduction of work in late-activated regions was primarily responsible for acutely improved myocardial energy efficiency, but these improvements after CRT were actually greater in long-term non-responders than in responders. Rather, the property of the work distribution that normalized after CRT and best predicted reverse ventricular remodeling was the size of the smaller region―primarily in the septum—having net systolic work done on it by the rest of the heart. Recently, Russell, et al. (7) tested a new method for estimating the “wasted work ratio” (the ratio of the sum of negative work to the sum of positive work performed by the myocardium); however, their method relied on simplified assumptions about regional myocardial mechanics that avoided the computation of regional wall stresses and predicted greatest wasted work on the RV free wall rather than in the septum.

Spatially heterogeneous systolic wall strains during dyssynchrony can contribute directly to work heterogeneity and are affected by CRT. Reversal by CRT of septal rebound stretch following premature shortening in the early-activated septum has been suggested as an indicator of reverse remodeling after resynchronization (38). In our patient-specific models at baseline (ventricular dyssynchrony), early-activated regions in the septum shortened early and stretched late during systole (**Fig. S1, Movie M1**). Tangney, et al (34) showed that late systolic stretches in papillary muscles, similar to those seen in early-activated regions of our patient-specific models, caused reduced or negative fiber work. Examining the magnitude of late septal systolic stretches in the baseline models, we saw a significant correlation with CRT outcomes (R^2^=0.48). However, there was a much weaker correlation between the decrease in septal rebound stretch after CRT and the 6-month reduction in end-systolic volume (R^2^=0.10). This suggests that while late systolic stretch may be an important component of negative septal work (**Fig. S2**), the therapeutic mechanism of CRT is more likely to be associated with its beneficial effects on regions of negative work than on septal systolic stretch *per se*.

### Work can be estimated with fewer clinical measurements compared to patient-specific modeling

To enable more widespread assessment in clinical CRT patients, we tested the accuracy of simplified approaches to estimate regional myocardial stress and strain using our computational models. The greater heterogeneity in work observed in CRT responders was also observed when work was estimated as the pressure-strain area (P_LHC_SA, P_gen_SA, P_gen,scaled_SA). We also evaluated the metrics found to be predictive in this cohort (negative work in the LV and septum). We found that approximations can be used to estimate the percentage of the LV surface performing negative work and accuracy is sufficient to maintain differences between CRT responders and non-responders. Our strain estimates were significantly simplified as values were derived from endocardial area change (which is being used in ECG-gated analysis of CT images). Compared to the computational estimates, surface assessment differs as it does not directly estimate the volume of the myocardium performing negative work. However, earlier comparison of regional strain estimated from endocardial surface area changes agreed with strain measured on CMR (18).

By testing different simplifications, we evaluated the need for patient-specific information when estimating myocardial stress. Somewhat surprisingly, P_LHC_SA had the most consistent (across metrics) agreement with patient-specific modeling work. We expected patient-specific pressure waveforms to perform better than generic waveforms as generic pressures likely overlook the hemodynamic timing differences that are incorporated in the catheter pressure. However, we would have expected that the addition of shape information would have improved agreement. This was not the case for WS_ED_SA or WS_TV_SA. Our use of ventricular pressure as a surrogate for stress is supported by earlier work which found agreement with regional oxygen uptake(10), accuracy across several physiologic states(39), and clinical utility(24,40,41). However, earlier work has also documented the limitation of pressure-strain estimates(42). In this work, our goal was not to replace stress-strain but to use the framework to identify a more readily available surrogate of regional myocardial work that can be applied to CRT patients.

### Limitations

Although the patient cohort in this study was small owing to the extensive and invasive measurements in severely diseased patients, the statistical findings were highly consistent. Four patients were echocardiographic responders and four were non-responders. Negative ventricular work volume fractions correlated most strongly with ΔESV_LV_, were significantly higher in responders at baseline, and decreased significantly more with CRT in responders. In addition, the long term reductions (and increases) in LV end-systolic volumes in the patient cohort spanned a wide range of values, which are usually seen in most DHF patients (43).

Although the baseline models were fit to a comprehensive, clinical functional data set, the CRT models only made use of known locations and interventricular delay times. Therefore, we used measurements of LV pressures and ECGs after the pacemaker was turned on as an independent test of the models. While peak pressures and pressure rates showed good agreement with measurements, model-predicted changes in QRS durations were not always consistent with measurements, possibly due to functional mechanisms such as lead capture delay, which were not included in the model. QRS lengthening is known to occur after CRT, and biventricular pacing can still provide clinical benefit in such cases (44).

RS_CT_ was initially designed to capture regional shortening from 4D cardiac CT data. In this study, subjects underwent 3D cardiac CT acquired at end-diastole, not ECG-gated cardiac CT. Therefore, RS_CT_ was quantified with temporal geometric changes informed by the patient-specific model. Additionally, myocardial work estimates WS_ED_SA and WS_TV_SA utilized shape information derived from the patient specific model, not from a direct clinical measurement. However, the patient-specific model yields physiological parameters that match clinical measurements (13). While we evaluated the importance of patient-specific peak pressures for a simple work estimate, we did not evaluate the importance of the timing of the pressure waveform in achieving an accurate work measurement. Given that the P_LHC_SA estimate had the strongest agreement with PSM-based work (**Supplement Simplified Work Estimates**), it is likely that both pressure values and timing are needed to accurately measure work. Our simplified myocardial work estimates were not evaluated in patients after CRT. Therefore, the ability for our work estimates to detect LV remodeling after CRT is left for future work.

### Conclusions

This study provides important new insights into reverse ventricular remodeling after CRT in patients with dyssynchronous heart failure. Patient-specific models confirmed in humans that CRT improves the mechanical efficiency of ventricular pumping and reduces the heterogeneity of regional myocardial work. However, neither the improvement in efficiency nor the decrease in heterogeneity *per se* explained the differences in the long-term reverse remodeling response between patients. While CRT decreased regions of high work in the late-activated regions and decreased regions of negative work in early-activated regions, it was only the latter that predicted the patient outcomes. This suggests that those reversible changes in perfusion, metabolism, and cell signaling that are seen primarily in the early-activated regions of the septum in patients with left bundle branch block may be important determinants of reverse ventricular remodeling after CRT. These findings suggest that CRT pacing protocols aimed at decreasing regions of negative myocardial work (rather than improving global work efficiency or decreasing regions of high work) may have greater long-term therapeutic benefit. Finally, our sensitivity analysis suggests that patient-specific models based on a minimal clinical data set including geometry, activation pattern, and systemic hemodynamics can make accurate estimates of regional work distributions and potentially support clinical decision-making for CRT candidates and predicting their outcomes. Further, simplified estimates of myocardial work based on limited clinical data were shown to stratify CRT responders from non-responders.

## Supporting information

Supplement Simplified Work Estimates

## Abbreviations

LBBB: Left Bundle-Branch Block
CRT: Cardiac Resynchronization Therapy
LV: Left Ventricle
ΔESV_LV_: Change in end-systolic LV volume 6 months after CRT
V_f_: Volume fraction
LVNW: LV Negative Work
STNW: Septal Negative Work

## Acknowledgments

We thank Jeff van Dorn for providing technical support in running the Continuity software. This work was supported by NIH grants F31HL165881 (to A.C), K01HL143113 (to F.C.), 1R01HL96544 (to A.D.M.), 1R01HL105242 (to A.D.M.), 1R01HL121754 (to A.D.M.), 8P41GM103426 (to R. Amaro and A.D.M.), P50GM094503 (to D. Beard and A.D.M.), an Additional Ventures Expansion Award, and the Saving Tiny Hearts Society.

## Author Contributions

D.E.K., S.M.N., J.H.O., R.C.P.K., and A.D.M. conceived and designed the study; D.E.K. recruited and consented patients and performed the ICD implantation procedure; A.C., A.K., C.V., K.V., developed and executed the computational models; A.C., A.J., F.C., A.D.M. designed the simplified modeling study and analysis; R.C.P.K. developed the methods for computational modeling; all authors contributed to interpretation of data and results; A.C., A.K., C.V., D.E.K., F.C., A.D.M. wrote the manuscript; and all authors provided edits; reviewed and approved the manuscript.

## Data and Materials Availability

Continuity 6.4 was used to run the simulations and the software can be downloaded from http://www.continuity.ucsd.edu/. Deidentified patient-specific models will be made available upon acceptance for publication using Dryad (which assigns a unique DOI for each dataset) and a Github repository will be used to share research code.

