## Supplemental Methods for "Successful Cardiac Resynchronization Therapy Reduces Negative Septal Work in Patient-Specific Models of Dyssynchronous Heart Failure"

#### Clinical Study

Several types of imaging data sets were obtained before device implantation. End-diastolic ventricular geometry was imaged by 64-slice computed tomography (CT) (Lightspeed VCG, GE Medical Systems, Milwaukee, WI) with an isotropic in-plane resolution of 0.49 mm (512×512 pixels) and slice thickness of 0.625 mm. In two patients, end-diastolic geometry was imaged by magnetic resonance (MR) imaging (specifications) with an in-plane resolution of 0.9 mm (128×128 pixels) and slice thickness of 2 mm. In five patients with myocardial infarction, affected regions were localized by singe-photon emission computed tomography (SPECT) (technetium sestamibi, ECAM-2, Siemens Medical Solutions, Hoffman Estates, IL) during rest and stress with an isotropic in-plane resolution of 6.59 mm and 10 mm slice thickness. Table 1 lists the locations of the infarcts.

Electrophysiology and hemodynamic measurements were obtained before and after device implantation. Standard 12-lead body-surface electrocardiograms (ECGs) were recorded (Bard Pro, Bard Electrophysiology, Lowell, MA) over 5-6 beats at 1 kHz sampling rate. Vectorcardiograms (VCGs) were derived from the ECGs (Kors, 1990). Left and right ventricular cavity pressures were continuously monitored by cardiac catheterization over several beats at 1 kHz sampling rate. Mean aortic pressure was recorded by sphygmomanometer (Bard Pro) measurements. Two-dimensional (2D) transthoracic echocardiography (TTE) (Sonos, Philips Medical IE33, Bothell, WA) recorded two- and four-chamber long-axis views of the ventricles over two beats. End-diastolic LV volumes and valve diameters (aortic and mitral annuli) were computed from the images. Continuous-wave Doppler TTE estimated regurgitant and forward stroke volumes by the velocity time integral (VTI) of hemodynamic flows through the mitral and aortic valves. **Table 1** summarizes patient characteristics at baseline.

Pacing devices were implanted with pace/sense leads in the right atrial appendage (RA lead), RV apical septum (RV lead), and the LV free wall (lead V) epicardial surface through a branch of the coronary sinus. One patient (BiV6) had a previously implanted RV pacemaker. Final LV lead positions on the ventricles were localized from bi-plane chest x-ray images by an expert electrophysiologist (DEK). A-V delays were programmed to 161.25 ± 21ms (paced) and 121.25 ± 14 ms (sensed). The V-V delay, (the delay between LV and RV lead pacing) was programmed to 19.4 ± 16.6ms according to an algorithm (QuickOpt, St. Jude) to maximize cardiac output from Doppler TTE. Following 3 ± 1 months of bi-ventricular pacing, the V-V delays were reprogrammed to 27.5 ± 17.9ms to maintain maximum stroke volume by transthoracic echocardiography.

#### Patient Specific Cardiovascular Model

#### Anatomy

Biventricular anatomical model construction from clinical and empirical data have been described previously (1,2). Briefly, CT/MR images of the ventricles at end-diastole were segmented and triangulated. A high-order cubic Hermite biventricular mesh atlas with 64 hexahedral elements was fitted to the triangulation using an iterative linear least squares algorithm (3). The resulting mesh was refined to 128 elements for biomechanics simulations. A coarser mesh atlas with 132 linear hexahedral elements was co-registered to its parent, converted to cubic Hermite hexahedrals (4), and refined to 8448 elements for electrophysiology simulations (5).

Human ventricular fiber architecture was modeled using a previous reconstruction fitted to diffusion tensor MR (DT-MR) images of an explanted human donor heart (6). The fiber architecture of the explanted heart was mapped to each patient-specific geometry via large deformation diffeomorphic mapping and tensor reorientation, to account for changes in the fiber structure due to patient-specific differences in the ventricular shape. Local orthogonal fiber, sheet, and sheet-normal axes were computed from the eigenvectors of the fitted tensors.

Regions of myocardial infarction were localized from rest and stress SPECT by an expert cardiac electrophysiologist and nuclear radiologist. Regions of depressed perfusion were manually demarcated on the CT/MR triangulations and registered in each patient-specific mesh as binary fields of normal and infarcted tissue.

#### Electrophysiology

Action potential propagation was modeled as mono-domain reaction-diffusion with human ventricular myocyte ionic currents (7). Ion channel properties (e.g. K^+^ current, Na-Ca exchanger) were adjusted to approximate observed changes in AP morphology during HF (8). Patient-specific 3D regional depolarization times at baseline were estimated by optimizing myocardial electrical conductivities and an ectopic stimulus located in the RV sub-endocardium to match vectorcardiograms derived from measured ECGs (5). Using the optimized baseline electrophysiology properties, acute CRT depolarization patterns for paced beats were simulated by applying stimuli according the prescribed V-V lead delays and locations in the anatomical model.

#### Biomechanics

The unloaded ventricular geometry at zero pressure and active and passive biomechanics properties at baseline were estimated as described in previous work (2). Using an empirical PV relation (9) and transversely isotropic passive stiffness (10), the diastolic pressure-volume (PV) loading curves for the unloaded ventricles were matched to the observed end-diastolic pressure and volume measured by intra-cardiac catheterization and echocardiography. The estimated unloaded geometry served as the reference configuration for subsequent stress, strain computations. Left and right systolic pressure traces were matched to the observed pressure traces (particularly peak systolic pressure, dP/dt_max_, and dP/dt_min_) and end-systolic volume by adjusting parameters of an active contraction model (11). Local onset of contraction was defined by the estimated local depolarization times in the electrophysiology model.

#### Hemodynamics

A closed-loop, self-adaptive model of the systemic and pulmonary circulation (CircAdapt, (12)) was coupled to the biomechanics model via pressure boundary conditions to compute systolic and diastolic pressure traces (1,2). Measured valve annuli diameters from 2D TTE were input directly as dimensional parameters, while others (e.g. atrial diameter) were predetermined within normal human physiologic limits. Aortic and pulmonary arterial length parameters were manually adjusted to match measured systolic pressures. Remaining circulatory parameters (e.g. vessel compliance) were self-adapted during non-steady state beats to match the cardiac output (accounting for mitral regurgitation) measured from Doppler TTE and aortic pressure measured by sphygmomanometer. The fully parameterized hemodynamic model was simulated over 11 beats at measured heart rates until steady-state pressure traces were obtained.

#### Global and Regional Myocardial Work Metrics

Principal (fiber, sheet, and sheet-normal) myocardial stresses and strains (referenced to the unloaded geometry) were computed during the 10th steady-state beat at the Gauss-Legendre quadrature points (27 per element, 3,456 total) of the biomechanics meshes.

#### Sensitivity Analysis

The patient-averaged geometry (-G) was computed from the spatial average of the patient-specific geometries. The patient-averaged LBBB activation pattern (-A) was simulated using the spatial average of the ectopic stimulus location in the RV and myocardial conductivities in the patient-averaged geometry. Patient-averaged CRT activation pattern was simulated using the average of the V-lead locations and V-V delays. Patient-averaged biomechanics (-M) and hemodynamics (-H) were obtained by averaging the optimized parameters of the biomechanics and the hemodynamics models, respectively. The patient-averaged unloaded geometry was re-estimated from the averaged biomechanics parameters. Regions of myocardial infarction were removed entirely in ischemic patients (-I). Finally, the simultaneous interaction effect of patient-specific geometry and LBBB activation pattern (-GA) was evaluated.

### Supplemental Figures

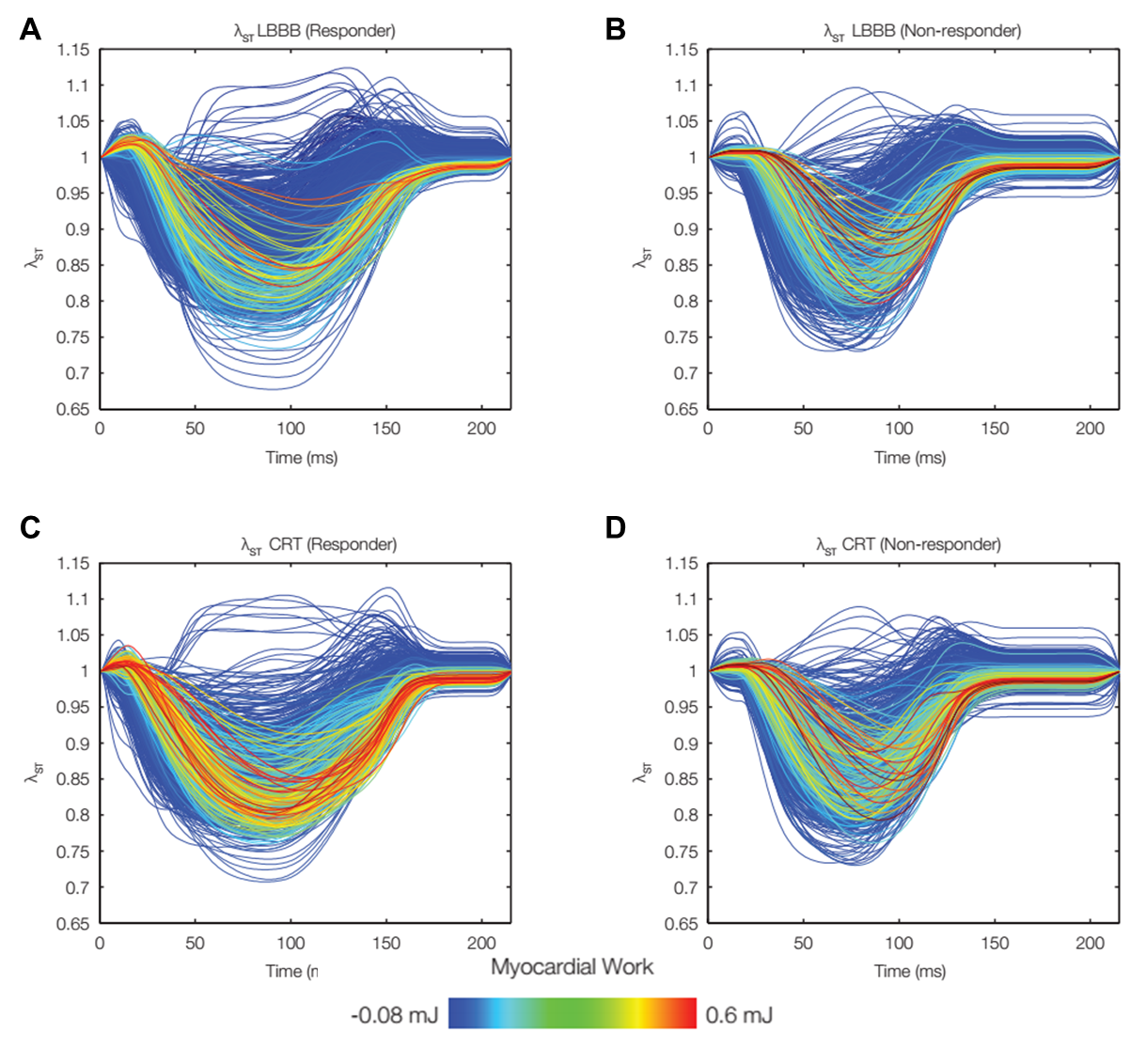

**Fig. S1: Late systolic stretch in the septum during LBBB and CRT.** Time courses of fiber stretch ratio referred to end-diastole at model integration points in the septum during LBBB (A and B) and CRT (C and D) in one responder (BiV6, A and C) and one non-responder (BiV7, B and D). The lines are colored according to the local myocardial work density. At baseline, late septal stretch (λ_ST_>1.0) and negative myocardial work (blue) are visibly more prevalent in a responder (A) than in a non-responder (B). During CRT, late stretch is reduced and work is increased for both responder (C) and non-responder (D).

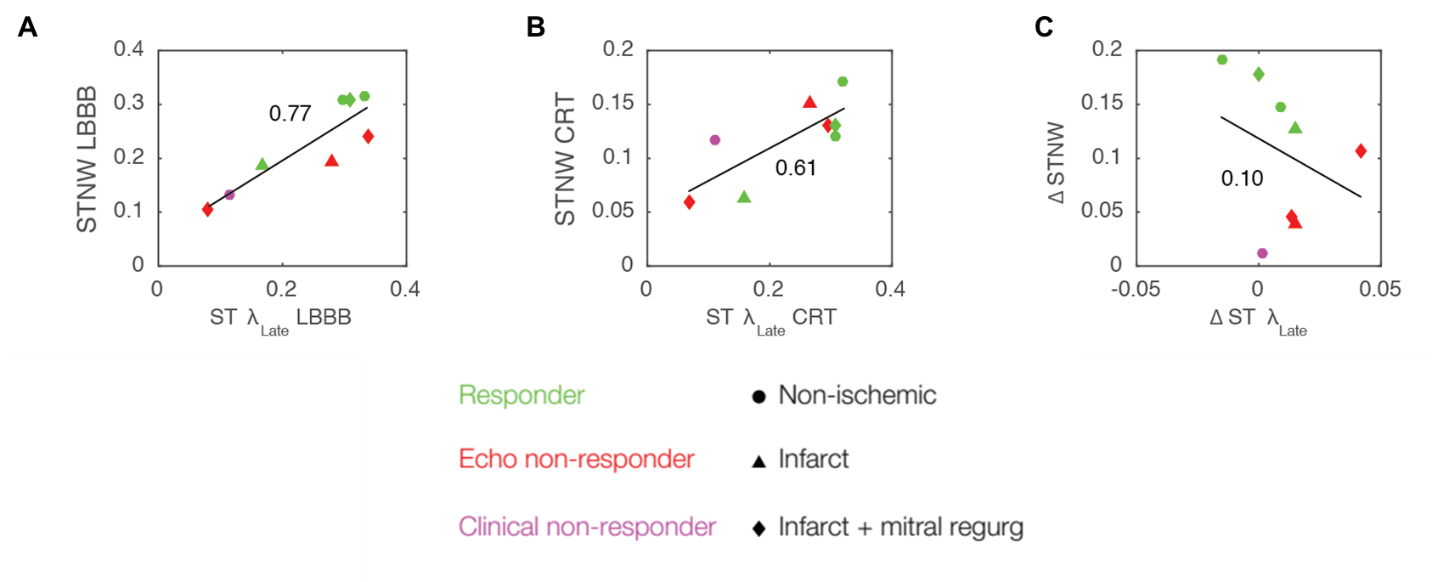

Fig. S2: Volume fraction of the septum with late systolic stretch correlates with the volume fraction of the septum performing negative work in both LBBB (A) and CRT (B). However, the change in the volume fraction of the septum with late stretch does not correlate with the change in negative work (C) nor with CRT outcomes as measured by reduction in LVESV (R^2^=0.1, data not shown). These findings suggest that late systolic stretch may be a useful marker for negative myocardial work, but that reducing late systolic stretch is not a therapeutic mechanism of CRT.

### Supplemental Movies

<BiV2 Echo Comparison LBBB CRT>

Movie M1. Early rapid abnormal motion of the septum (septal flash) is prominent at LBBB and is reversed in CRT in both echocardiographic imaging and in the patient-specific models. The movie shows the patient-specific model (in red) overlaid on the echocardiographic imaging of the patient (in black-and-white). Both the patient-specific model and the echocardiographic imaging show abnormal movement (marked with a green box) of the septum early due to earlier activation of the septal region compared to the LV. CRT activation reverses this abnormal motion of the septum, since the contraction is more synchronous.
