## Supplement Simplified Work Estimates for "Successful Cardiac Resynchronization Therapy Reduces Negative Septal Work in Patient-Specific Models of Dyssynchronous Heart Failure"

**Supplementary Material on Simplified Estimates of Myocardial Work**

**Methods**

Five simplified myocardial work estimates were evaluated to assess the extent of patient-specific information needed to measure segmental LV work. Estimates were calculated as the area enclosed by approximations of stress-strain loops. These approximations include 1) left heart catheterization derived pressure-strain area (P_LHC_SA), 2) end-diastolic Laplace wall stress-strain area (WS_ED_SA), 3) time-varying Laplace wall stress-strain area (WS_TV_SA), 4) generic LV pressure-strain area (P_generic_SA), and 5) generic pressure-strain area with pressure scaled to patient-specific peak pressure (P_generic,scaled_SA). The calculations for the simplified strain and myocardial work estimates are described in the *Simplified Myocardial Work Estimates* section of the methods in the manuscript. To assess agreement between myocardial strain and work measured with simple estimates and patient-specific modeling, each measurement was averaged for all 17 AHA segments to achieve one strain curve and one myocardial work measurement per segment.

*Statistical Analysis*

Simple work estimates were correlated with work measured with the patient-specific model using simple linear regression. These estimates were determined to not be normally distributed by the Shapiro-Wilk test, therefore significance was determined by Spearman correlation.

**Results**

*Simplified regional strain agrees strongly with regional strain computed with patient-specific modeling*

Segmental strain computed with RS_CT_ correlated strongly to segmental strain computed with patient-specific modeling across all patients (median: R = 0.76, IQR: 0.71 – 0.81, **Table S1**). This strong correlation held when including all segmental strain measurements for all patients (R = 0.66, p < 0.01, **Fig. S1**).

*Simplified estimates of regional myocardial work agree with regional myocardial work computed from patient-specific models*

Spearman correlation coefficients of each comparison to patient-specific modeling results for each patient ranged from moderate to strong (**Table S2**). P_LHC_SA correlated strongly with the model calculation across patients (median R = 0.80, IQR: 0.73 – 0.82), and correlated significantly in patients BiV1-7. When including all segmental myocardial work measurements for all patients, P_LHC_SA significantly correlated to the model calculation (R = 0.77, p < 0.01, **Fig. S2A**).

Incorporating regional shape information into WS_ED_SA correlated moderately with the model calculation across patients (R = 0.68, IQR: 0.46 – 0.77), and correlated significantly in BiV1-6. WE_TV_SA also moderately correlated with the model-based myocardial work across patients (median R = 0.70, IQR: 0.61 – 0.73), with significant correlations for patients BiV1-7. Both the static and time-varying estimates of all LV segments in all patients correlated significantly with all segmental work measurements from the model (static: R = 0.71, p < 0.01, **Fig. S2B**; time-varying: R = 0.72, p < 0.01, **Fig. S2C**).

Use of a generic LV pressure curve in P_gen_SA correlated moderately across patients (median: R = 0.55, IQR: 0.38 – 0.76). When scaled to patient-specific peak pressures for P_gen,scaled_SA, the correlation became/remained moderate (median: R = 0.55, IQR: 0.38 – 0.76). In both cases, results in patients BiV1, 3, 4, 5, and 7 correlated significantly. When all LV segments from all patients were included, both estimates were moderately correlated to model-based segmental myocardial work (generic: R = 0.55, p < 0.01, **Fig. S2D**; scaled to peak pressure: R = 0.63, p < 0.01, **Fig. S2E**). However, Fisher’s z-transformation demonstrated the correlations were not found to be significantly different from one another (p > 0.05).

**Discussion**

*Work can be estimated with fewer clinical measurements compared to patient-specific modeling*

While myocardial work is clinically estimated as the ventricular pressure-strain loop area (PSA), multi-scale patient-modeling can measure work as the myocardial stress-strain loop area. Simple estimates of work derived from limited clinical information demonstrated good agreement with work measured with patient-specific modeling. P_LHC_SA had the strongest agreement with modeling-derived work. P_gen_SA had the weakest, yet moderate correlation (R = 0.55), which suggests the expected shape of the LV pressure waveform is important for measuring myocardial work. Incorporating additional patient-specific information improved agreement, which is demonstrated by P_gen,scaled_SA and P_LHC_SA. Sensitivity analysis of the model showed that work is highly sensitive to patient-specific hemodynamics, which supports our findings that P_LHC_SA, the work estimate with the most patient-specific LV hemodynamics, has the strongest correlation to the patient-specific modeling work measurement.

However, work estimates including local shape information improved agreement in only three patients: BiV3, 6, and 8. Laplace wall stress has been described as a simplified approximation of global mean wall stress (1,2). We anticipated this mean stress approximation to be advantageous for work estimates WS_ED_SA and WS_TV_SA, where segmental wall stress was estimated with mean segmental measures of LV wall thickness, endocardial radius, and endocardial strain. Despite our efforts to capture local shape information, it is likely that our measures of segmental wall thickness and radius overgeneralized spatial heterogeneities in LV shape, thus leading to a worse work estimate. Given the clinical challenges in capturing regional LV shape, it is possible that measuring regional wall stress is difficult without intensive finite element modeling of LV mechanics, as demonstrated by Gsell et al. (3).

Work estimates for BiV8 did not correlate to work measured with PSM. However, our peak strain estimate was strongly correlated (R = 0.71) to strain measured with PSM, indicating that our stress approximations failed to reflect the PSM-informed stress.

**Do we need additional discussion here?**

**Conclusion**

Left ventricular myocardial work and strain estimated with limited clinical measurements agrees with LV performance computed with patient-specific modeling. Therefore, a more readily available estimate of LV performance, especially when computed with patient-specific strain and hemodynamics, can serve as a surrogate of regional myocardial work that can inform clinical decision making in CRT patients.

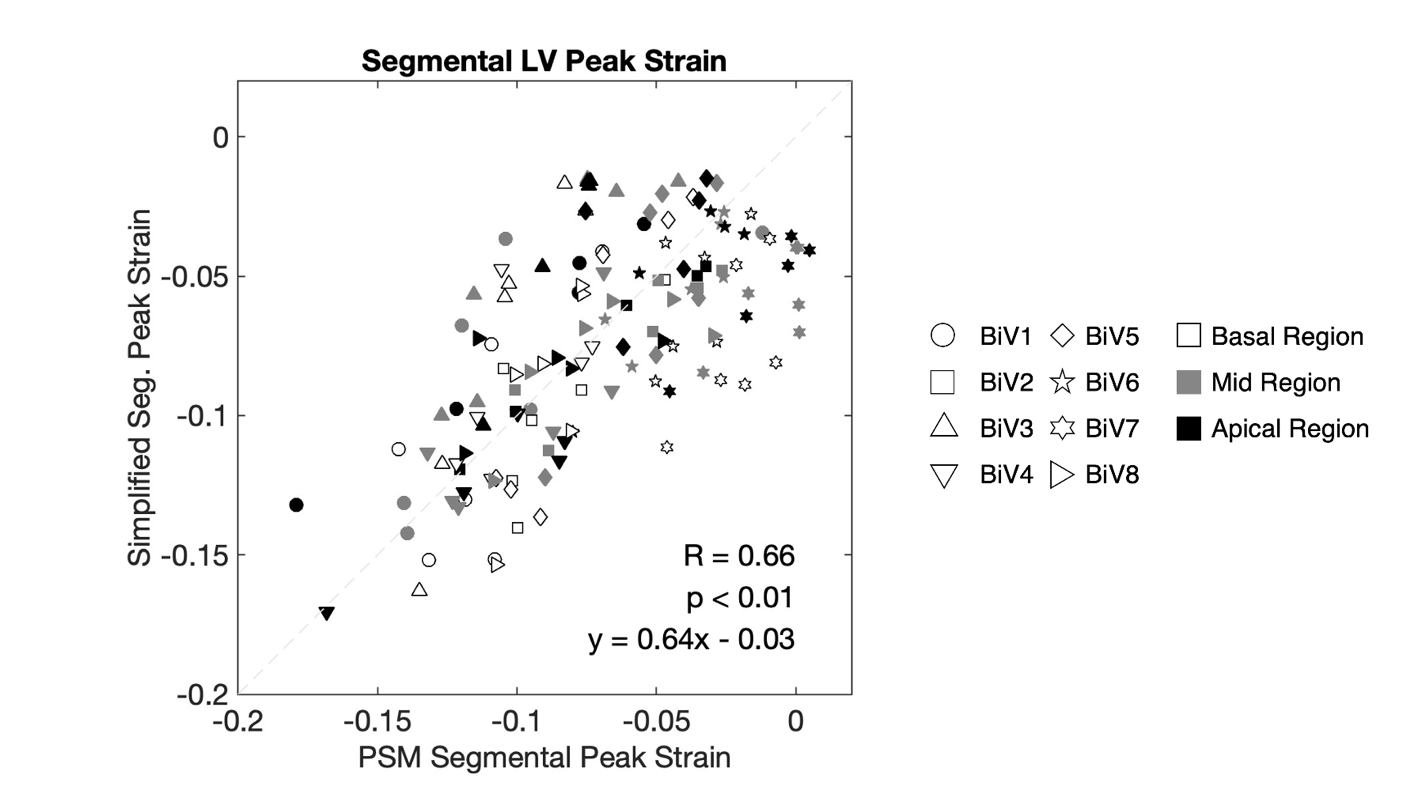

**Fig. S1: Strain estimated from endocardial surface area change correlates strongly with PSM derived strain.** Spearman correlation of peak segmental LV strain measured with endocardial surface tracking and peak segmental LV strain measured with patient-specific modeling. Each datapoint represents one segment of the 17-segment AHA model of one subject. Shapes denote work measured in each patient and described above. Shading color denotes work measured in the basal (black), mid (gray), and apical (white) regions of the LV. The dashed line is the identity line. Evaluating segmental work measurements for one subject demonstrates that the relationship between CT- and PSM-derived work measurements is independent of segment location.

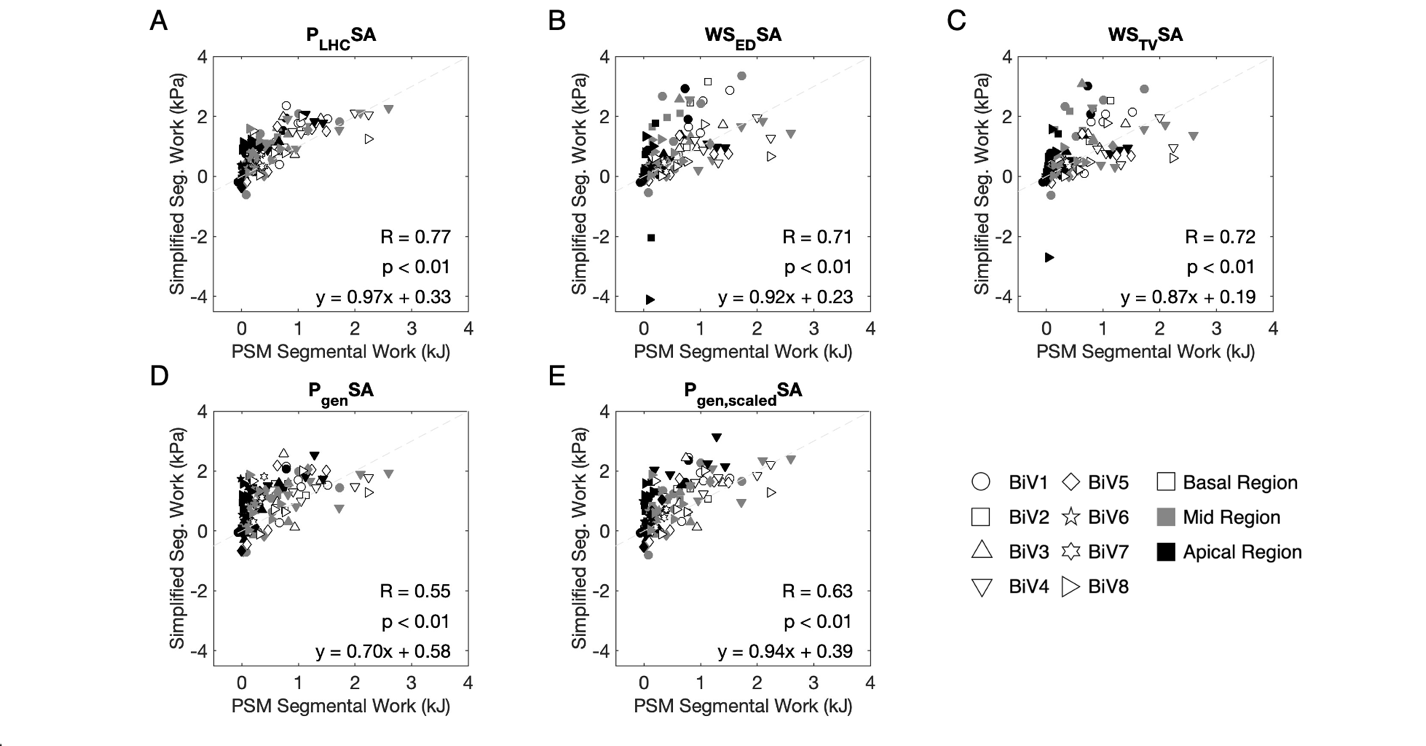

Fig. S2: Simplified estimates of myocardial work correlate with myocardial work measured with patient-specific modeling for all patients. Spearman correlations between segmental LV work measured as the left heart catheterization (LHC) LV pressure-strain loop area (P_LHC_SA, A), end-diastolic wall stress-strain loop area (WS_ED_SA, B), time-varying wall stress-strain loop area (WS_TV_SA, C) generic LV pressure-strain loop area (P_gen_SA, D), generic LV pressure scaled to patient peak pressure-strain loop area (P_gen,scaled_SA, E), and myocardial work measured with patient-specific modeling are shown. Each datapoint represents one segment of the 17-segment AHA model of one subject. Shapes denote work measured in each patient and described above. Shading color denotes work measured in the basal (black), mid (gray), and apical (white) regions of the LV. The dashed line is the identity line. P_LHC_SA has the strongest correlation to the control. Correlations between simpler myocardial work estimates and work calculated with the patient-specific model range from moderate to strong.

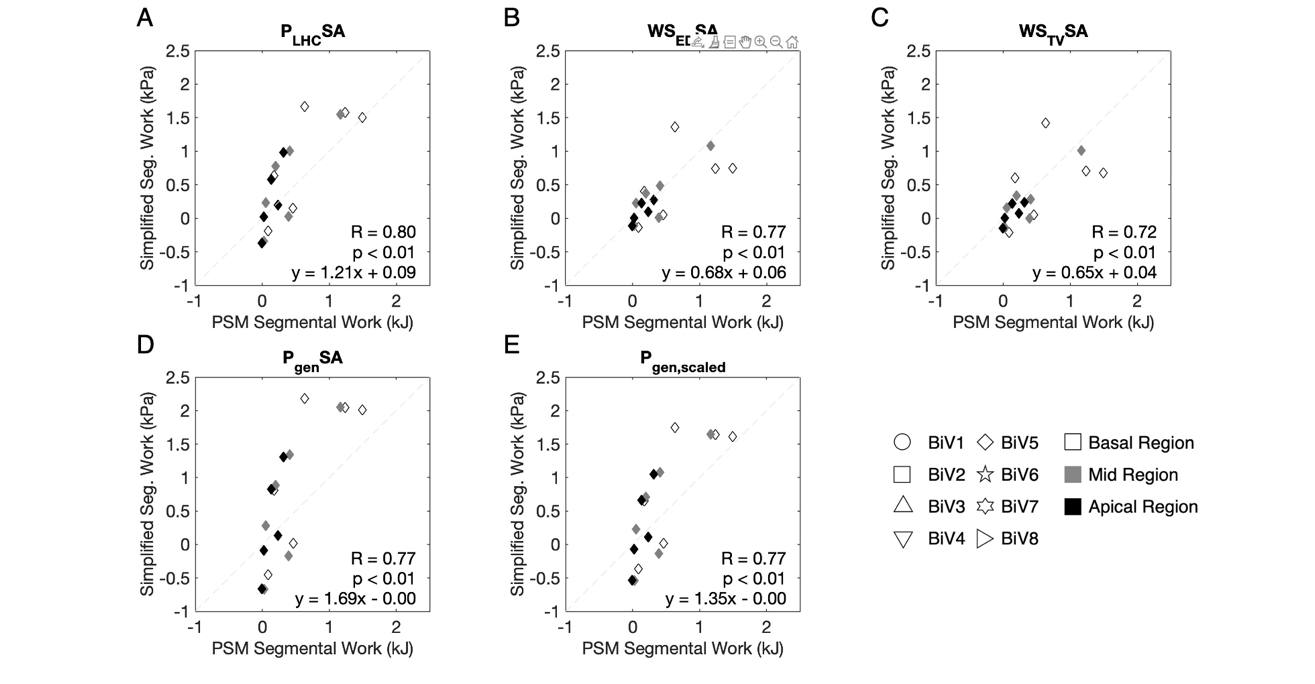

Fig. S3: Simplified estimates of myocardial work correlate with myocardial work measured with patient-specific modeling for a representative patient. Spearman correlations between segmental LV work measured as the left heart catheterization (LHC) LV pressure-strain loop area (P_LHC_SA, A), end-diastolic wall stress-strain loop area (WS_ED_SA, B), time-varying wall stress-strain loop area (WS_TV_SA, C) generic LV pressure-strain loop area (P_gen_SA, D), generic LV pressure scaled to patient peak pressure-strain loop area (P_gen,scaled_SA, E), and myocardial work measured with patient-specific modeling are shown. Each datapoint represents one segment of the 17-segment AHA model of one subject. Shapes denote work measured in each patient and described above. Shading color denotes work measured in the basal (black), mid (gray), and apical (white) regions of the LV. The dashed line is the identity line. Evaluating segmental work measurements for one subject demonstrates that the relationship between CT- and PSM-derived work measurements is independent of segment location.

**Table S1: Summary of simplified regional strain compared to PSM regional strain.**

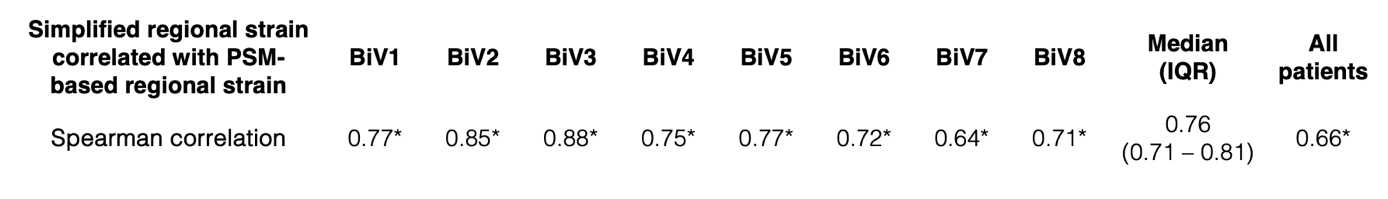

Spearman correlations between segmental LV strain estimated with endocardial tracking on CT and segmental strain measured with patient-specific modeling. Correlations for each patient ranged from strong-to-very strong. The correlation of segmental strain measured across all patients was also strong; this relationship is displayed in **Fig. S1**.

**Table S2: Summary of simple myocardial work estimates compared to PSM measurements.**

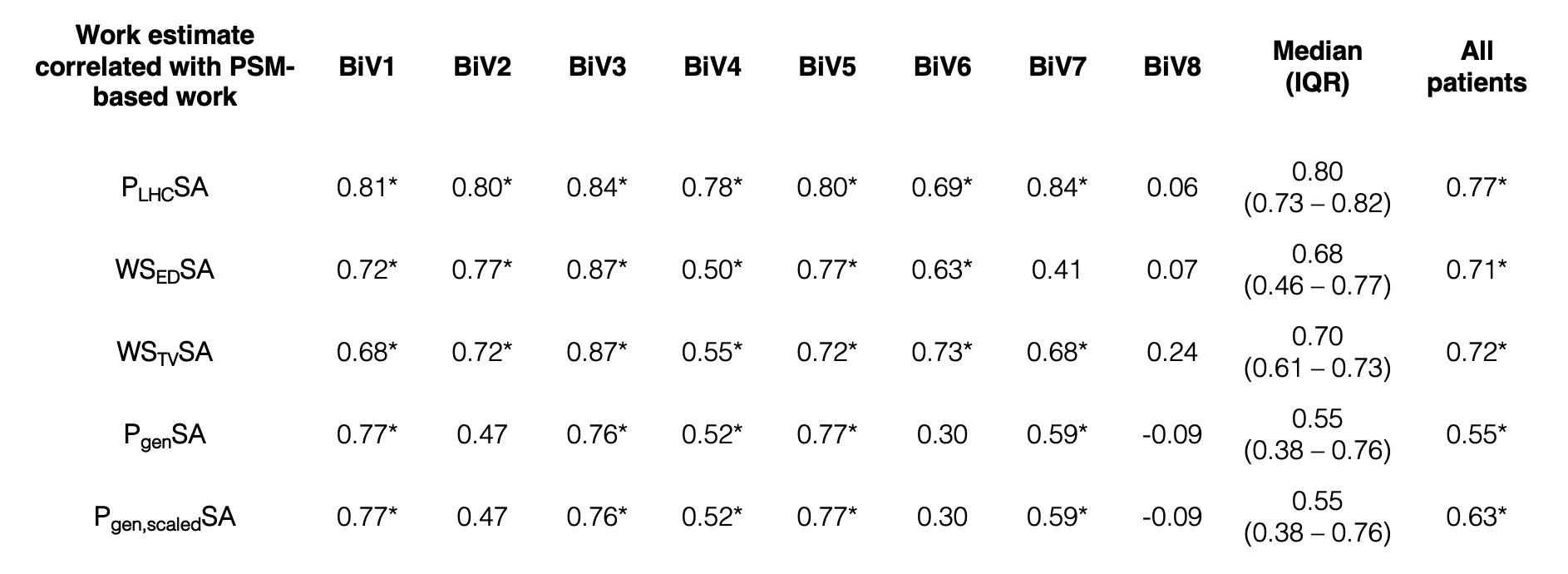

Spearman correlations between segmental LV work measured as the left heart catheterization (LHC) LV pressure-strain loop area (P_LHC_SA), end-diastolic wall stress-strain loop area (WS_ED_SA), time-varying wall stress-strain loop area (WS_TV_SA) generic LV pressure-strain loop area (P_gen_SA), generic LV pressure scaled to patient peak pressure-strain loop area (P_gen,scaled_SA), and myocardial work measured with patient-specific modeling are shown. Significant correlations ranged from moderate-to-very strong across work estimates and patients. No myocardial work estimates for BiV8 correlated strongly with the control. Incorporating LV shape improved myocardial work correlations in BiV3, 6, and 8, but worsened in BiV1, 2, 4, 5, and 7. Myocardial work estimated as the left heart catheterization (LHC) LV pressure-strain loop area had the strongest correlation with the control when considering all patients. Correlation between work measurements across all patients (right most column) are displayed in Fig. S2-S3.

*indicates statistical significance (p<0.05).
